# Dual-Chassis Strategy for Bridging Adaptive Evolution and Rational Design for Synthetic Biology

**DOI:** 10.64898/2026.06.02.729570

**Authors:** Kesi Kurnia, Isaac Gifford, Ville Santala, Jeffrey E. Barrick, Suvi Santala

**Affiliations:** Faculty of Engineering and Natural Sciences, Tampere University, Tampere, 33720, Finland; Department of Molecular Biosciences, Center for Systems and Synthetic Biology, The University of Texas at Austin, Austin, TX 78712, United States; Department of Microbiology, Genetics, and Immunology, Michigan State University, MI 48824, United States

## Abstract

Genome streamlining and pathway refactoring are powerful strategies for constructing controllable microbial chassis for both fundamental studies and applications. While rational design benefits from reduced genetic complexity, adaptive laboratory evolution (ALE) thrives on metabolic redundancy, creating a mismatch between optimal hosts for design and evolution. Here, we introduce a dual chassis framework (DUET) in which rational pathway construction and adaptive evolution are first carried out in an evolution-competent host, and the resulting optimized designs are subsequently transferred into a genetically stable chassis for deployment. Using the naturally evolvable bacterium *Acinetobacter baylyi* ADP1 and its genome-stabilized derivative (ISx), we applied this framework to the β-ketoadipate pathway, a central hub for aromatic compound catabolism. We first streamlined the native network by deleting individual pathway branches and then engineered a minimal synthetic route that merges protocatechuate and catechol metabolism. Subsequent ALE enabled efficient growth through this synthetic pathway, and reverse-engineering identified key adaptive mutations underlying functional recovery. Both the synthetic pathway and the mutations were transferred unchanged into ISx, where robust growth was maintained without further adaptation. These results demonstrate that DUET enables portable, host-independent deployment of rational metabolic streamlining combined with evolution, providing a generalizable strategy for building reduced yet robust microbial platforms.

**GRAPHICAL ABSTRACT:** 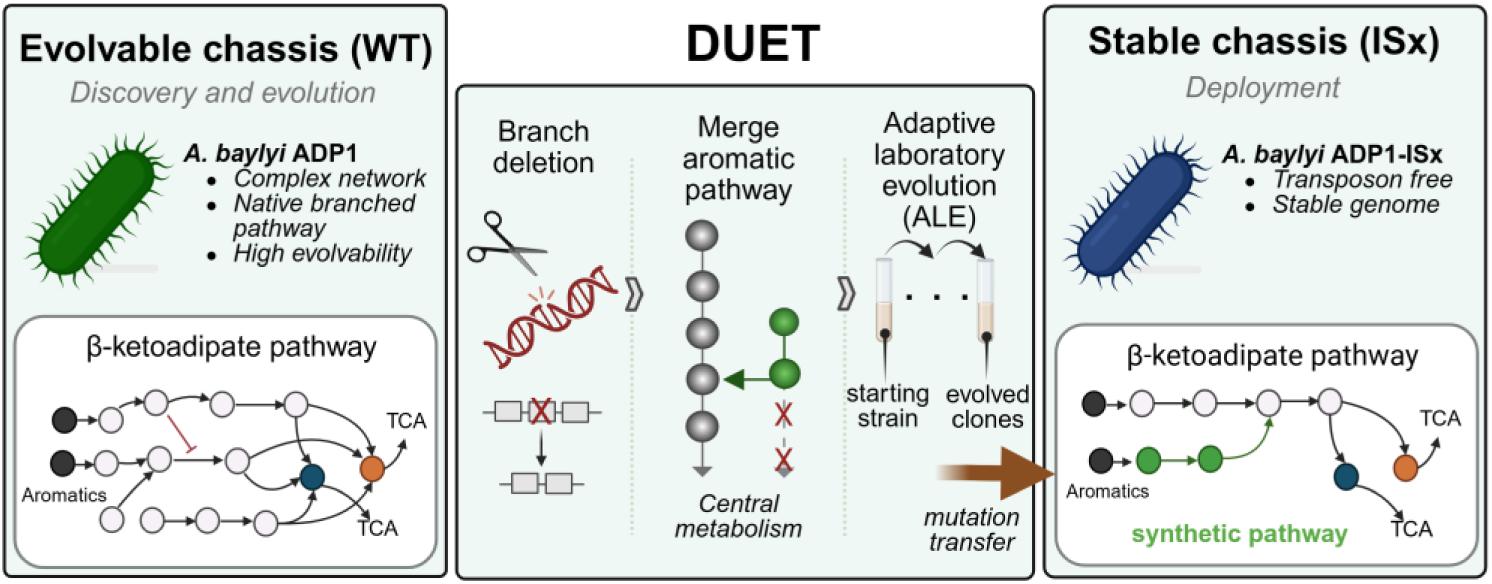

## INTRODUCTION

In synthetic biology, genome streamlining followed by refactoring of natural pathways offers powerful strategies for developing more controllable cellular platforms, both for fundamental research and biotechnological applications. By removing non-essential genes, mobile elements, and redundant functions, streamlining can improve performance under relevant conditions, enhance genome stability, and increased predictability by reducing native regulatory complexity (1, 2). For example, the minimized *Escherichia coli* strain MS56 was derived from MG1655 through a 23% genome reduction, yielding a robust chassis with enhanced genetic stability and improved recombinant protein production (3). Similarly, the streamlined derivatives *Pseudomonas putida* EM42 and EM383 were engineered through systematic removal of prophages, transposons, and other non-essential loci that compromise genetic stability or drain metabolic resources. As a result, these strains exhibit greater stability, improved heterologous gene expression, and reduced metabolic burden relative to the parental KT2440 strain (4). In parallel, pathway refactoring enables the redesign of genetic elements and regulatory architectures to decouple them from native control, thereby facilitating modularity and precise control of rationally designed synthetic pathways (5). Together, these strategies advance the development of standardized, programmable chassis that can be flexibly repurposed for metabolic engineering, pathway dissection, and industrial deployment.

Although refactored pathways are designed using state-of-the-art principles, they often underperform compared to their native counterparts. This discrepancy largely arises from host-dependent constraints, including regulatory crosstalk, resource competition, and cofactor or energy imbalances, which collectively reduce pathway activity and stability and give rise to context-dependent behaviour (6). To overcome these limitations, adaptive laboratory evolution (ALE) enables engineered hosts to acquire mutations that improve fitness and pathway performance under defined selective pressures (2, 7, 8). Rather than addressing these constraints through further rational modification, ALE leverages natural selection to reprogram global regulation and metabolism, allowing cells to adaptively resolve design-imposed imbalances. For example, in genome-reduced *E. coli* MS56, evolution restored near–wild-type growth through transcriptomic and translational reprogramming driven by mutations in global regulators such as *rpoD* and *rpoS* (9). Comparable adaptive rewiring was observed in a stable genome *Acinetobacter baylyi* ADP1-ISx derivatives, where mutations in genes encoding post-transcriptional regulators including *csrA*, *hfq*, and *rnd* compensated for large chromosomal deletions (8). Likewise, in *P. putida* KT2440, evolution refined enzyme activity and regulatory control within aromatic catabolic pathways by selecting mutations in *fghA*, *vanB*, and *PP_3493* (10). Collectively, these studies demonstrate that evolution can re-establish systems-level homeostasis and reveal non-intuitive yet effective genetic solutions to design-imposed limitations.

However, optimal host properties required for rational design and for evolutionary adaptation are not always well aligned. For example, streamlined strains are attractive for their stability and predictability (11), yet they often lack the redundancy and plasticity that enable diverse adaptive trajectories, limiting their capacity to benefit from ALE precisely when adaptability is most needed. This tension reflects a broader challenge in synthetic biology: how to combine the controllability of engineered systems with the adaptability inherent to living cells.

Owing to the metabolic versatility, natural competence, and efficient genetic manipulation, *A. baylyi* ADP1 (later on referred to as ADP1) has previously emerged as a model organism for genetic, metabolic, and evolutionary studies (12–14). Moreover, ADP1 has been explored for the production of various value-added compounds from lignin-derived molecules including wax esters (15), 1-alkenes (16), naringenin (17), mevalonate (18), and *cis,cis*-muconic acid (19). In particular, the natural competence has been directly exploited to create automated DNA assembly workflows, enabling high-throughput cloning and pathway construction without reliance on *E. coli* intermediates (20). The genomic tractability and the rapid adaptability under selective pressures has also made it a frequent choice as a platform for evolutionary and reverse engineering studies (21), supported by the previously established tools such as RAMSES (22), CEMENT (23), and EASy (24). In addition, an insertion sequence (IS) free derivative of ADP1 was previously constructed (11); the stabilized-genome strain designated as ISx exhibits enhanced genetic stability and transformability, but reduced evolvability (11). Notably, ISx was subsequently used to develop a Golden Transformation procedure for modular genome engineering and CRISPR-Lock tools that stabilize genomic deletions (25).

The contrasting properties of ADP1 and ISx make the pair uniquely suited for a novel dual chassis strategy presented here, termed as DUal-chassis system for combined Engineering and evoluTion (DUET): the wild type serves as an evolutionary platform in which adaptive mutations can arise, while ISx provides a streamlined, predictable, and stable background for implementing and evaluating these solutions. Together, these strains enable a systematic dissection of how evolutionary innovations can be captured in complex genomes and transplanted into stable-genome hosts. DUET concept reflects the stepwise process by which adaptive innovations discovered through evolution are systematically translated into a streamlined chassis for rational design. By bridging design and evolution, this strategy offers a generalizable route to integrate adaptability with precision, effectively decoupling refactoring and evolution phases to enable more efficient development of high-performing synthetic cellular platforms.

As a model for streamlining and refactoring, we chose the β-ketoadipate pathway (KAP) in ADP1, which functions as the central hub of aromatic catabolism. This pathway channels diverse substrates through two parallel entry routes, the protocatechuate and catechol branches, into a common intermediate that feeds the tricarboxylic acid (TCA) cycle (26). The protocatechuate branch primarily processes hydroxybenzoates and phenylpropanoids, while the catechol branch targets benzoates and related compounds. Although KAP is broadly conserved among soil bacteria, it is subject to tight regulation, with each branch induced only by its corresponding substrate, often resulting in hierarchical rather than simultaneous co-metabolism (27–29). For example, benzoate typically induces the catechol branch and is consumed before 4-hydroxybenzoate, which activates the protocatechuate branch (27, 28). Such sequential utilization constrains overall carbon flux, limiting efficiency when multiple substrates are present. Previous efforts have improved single-substrate utilization or boosted titers of specific products, but systematic strategies to bypass these regulatory hierarchies and streamline the pathway are still lacking. Addressing this gap is essential for both fundamental understanding of pathway dynamics and applied efforts in areas such as lignin valorization and bioremediation.

In this study, we implemented a dual-chassis system within a design–build–test–learn (DBTL) framework to systematically streamline and reconstruct the KAP pathway in ADP1. Taking advantage of ADP1’s genetic accessibility, we constructed branch-specific deletion strains to isolate the catechol and protocatechuate arms, thereby eliminating cross-regulatory interference. This modular architecture enabled two complementary strategies: (i) division-of-labor designs for mixed aromatic substrates, and (ii) rational reconstruction of a minimal synthetic catabolic pathway. The reconstructed pathway was subsequently optimized through adaptive laboratory evolution (ALE), yielding strains capable of growth on the target substrate with only minimal gene sets. Reverse engineering of evolved variants identified causal mutations, providing insights into principles of metabolic robustness and trade-offs in pathway optimization. We introduced the mutations obtained from ADP1 into the genome-stabilized ISx strain, thereby bridging the gap between evolutionary exploration and rational design. Together, this work establishes the ADP1-ISx dual chassis as a powerful model system for probing the design rules of streamlined metabolism.

## MATERIAL AND METHODS

### Bacterial strains, plasmids, and culture conditions

The bacterial strains and plasmids used in this study are described in Supplementary Table S1 and S2. *E. coli* strains were routinely cultivated at 37 °C, *A. baylyi* ADP1 and derived strains were incubated at 30 °C in low salt LB medium (1 g NaCl, 10 g tryptone and 5 g yeast extract per liter). Throughout this study, ADP1-WT (DSM 24193, DSMZ) refers to the previously ADP1(S) strain (30). For grown-on plates, 1.5% agar (w/v) was added. For growth experiments, ADP1 strains were cultivated in minimal salt medium (MSM) (per Liter; 3.88 g K_2_HPO_4_, 1.63 g NaH_2_PO_4_, 2.0 g (NH_4_)_2_SO_4_, pH 7.0). The MSM was supplemented with a trace elements solution (per Liter; 10 mg/L ethylenediaminetetraacetic acid (EDTA), 0.1 g/L MgCl_2_·6H_2_O 2 mg/L ZnSO_4_·7H_2_O, 1 mg/L CaCl_2_·2H_2_O, 5 mg/L FeSO_4_·7H_2_O, 0.2 mg/L, Na_2_MoO_4_·2H_2_O, 0.2 mg/L CuSO_4_·5H_2_O, 0.4 mg/L CoCl_2_·6H_2_O, 1 mg/L MnCl_2_·2H_2_O). Benzoate, 4-hydroxybenzoate (4HB), *p*-coumarate, *trans*-ferulate, vanillate, and protocatechuate (PCA) stock solutions were prepared in water with a concentration of 100 – 200 mM and titrated with 5 M KOH to pH 8.3–8.5. Adipate solution was prepared at concentration of 1 M (pH 7.0). When required, antibiotics kanamycin (Km) 50 µg/mL, chloramphenicol (Cm) 25 μg/mL, or gentamycin (Gm) 15 μg/mL or the nucleoside analogue 3′-azido-2′,3′-dideoxythymidine (AZT) 200 μg/mL were added to the medium. For N-(3-oxohexanoyl) homoserine lactone (AHL), a 100 mM stock solution was prepared in DMSO. All aromatic compounds were purchased from Tokyo Chemical Industry (Japan), except for *trans*-ferulate and AHL which were purchased from Sigma-Aldrich (USA).

### General DNA manipulations and plasmid construction

DNA spacers and oligonucleotides used in this study are listed in Supplementary Table S3 and S4. Oligonucleotide primers were synthesized by Thermo Scientific (USA). DNA spacers were ordered from custom oligo synthesis Genescript Biotech (China). Commercial kits and enzymes were used according to the recommendations given by the corresponding manufacturers. Plasmid DNA, PCR amplicons, and gel fragments were purified using GeneJet plasmid miniprep, GeneJet PCR purification, or GeneJet gel extraction kits (Thermo Fisher Scientific). PCR amplifications were performed using Phusion HotStart II high-fidelity DNA polymerase (Thermo Fisher Scientific). The OneTaq master mix (New England Biolabs, USA) was used to perform colony PCRs. The BsaI, BsmBI, and T4 ligase enzymes (New England Biolabs) were used to carry out golden gate cloning. Genomic DNA was extracted using the Monarch Genomic DNA Purification Kit (New England Biolabs) or the GeneJET Genomic DNA Purification Kit (Thermo Fisher Scientific).

### Construction of gene deletion strains

To systematically disrupt aromatic and dicarboxylic acid degradation pathways in *A. baylyi* ADP1, four deletion strains were constructed. The catechol pathway deletion strain (ΔCAT) was generated by removing *antABC* (ACIAD2669–2671) and the *sal-are-ben-cat* cluster (ACIAD1424–1451). The protocatechuate pathway deletion strain (ΔPCA) was constructed by deleting the *van* operon (ACIAD0978–0983) and the *pca-qui-pob-hca* cluster (ACIAD1702–1728). The dicarboxylic acid branch deletion strain (ΔDCA) targeted *mucK* and the *dca* cluster (ACIAD1681–1699). A combined strain incorporating all deletions was designated as strain ΔKAP. The deleted genomic regions and their associated metabolic functions are shown in Supplementary Fig. S1 and Supplementary Table S5.

All gene deletions were carried out using the Golden Transformation technique, as previously described by Suárez et al., 2020 (25). Deletion constructs were assembled via Golden Gate Assembly (GGA) using BsaI, by combining ∼1-kb upstream (HR1) and downstream (HR2) homology arms with the pBTK622 plasmid carrying the *tdk-kanR* selection cassette. The assembled constructs were introduced into ADP1 by natural transformation, and transformants were selected on LB agar containing kanamycin. When transformation efficiency was low, alternative methods such as Puddle or Acinetobacter Minimum Succinate (ABMS) transformation (25) were employed using freshly assembled GGA products.

To remove the *tdk-kanR* and generate a markerless deletion, rescue cassettes (∼2 kb) were assembled using BsmBI and introduced to *tdk-kanR* strains. Selection was performed on LB plates supplemented AZT. Successful deletions were verified by colony PCR and confirmed by loss of kanamycin resistance. A schematic representation of constructing deletion strains of ADP1 is presented in Supplementary Figure S2.

### Growth characterization in 96 well-plates

ADP1 and derived strains were pre-cultured in MSM supplemented with 0.2% (w/v) casamino acids and 1 mM of the respective aromatic compound at 30 °C and 300 rpm overnight. Cells were washed twice with MSM without any carbon source by centrifuging the culture at 6000 x g for 6 minutes, discarding the supernatant, and resuspending the cells with MSM. The cells were then inoculated into 96-well plates containing 200 µL of MSM with the test aromatic compound as the sole carbon source at the concentrations listed in Supplementary Table S6. Cultures were prepared in triplicate with an initial OD_600_ of 0.05. Plates were incubated at 30 °C for 48 h, and OD_600_ was recorded hourly using a Spark multimode microplate reader (Tecan, Switzerland). Cultures were mixed by double orbital shaking at 6 mm amplitude and 54 rpm, twice per hour. OD_600_ values were baseline-corrected by subtracting the absorbance of cell-free medium wells. The maximum specific growth rate (μ_max_) was calculated during the exponential growth phase by calculating the slope of a linear regression fitted to the natural logarithm of average OD_600_ values over time. The lag phase was defined as the period from inoculation (starting OD_600_ = 0.05) until OD_600_ reached 0.08.

### CRISPR-Lock and co-culture experiments for mixed substrate utilization

The *tdk-kanR* module used to replace the native CRISPR array of ADP1 was amplified from the CRISPR-Ready strain (25), and introduced into ADP1 strains by natural transformation. Colonies exhibiting kanamycin resistance, indicating successful integration, were selected for further modification. Self-targeting spacer cassettes, containing 32 bp of the deleted region adjacent to a 5′-CC protospacer-adjacent motif (PAM) and flanking sequences, were assembled using BsmBI-based Golden Gate Assembly. These cassettes were introduced into the CRISPR-Ready variant of ADP1 strains, and transformants were selected on AZT-containing medium. Positive clones were verified by colony PCR and kanamycin sensitivity screening.

The resulting CRISPR-Lock strains were used in co-culture experiments to assess substrate consumption. Cultures were grown in 100-mL baffled flasks containing 20 mL of MSM supplemented with 4HB and benzoate. In brief, glycerol stocks were streaked onto LB agar and incubated overnight at 30 °C. Single colonies (n = 3) were inoculated into MSM containing 0.2% (w/v) casamino acids and 1 mM of each aromatic compound and cultured overnight at 30 °C with shaking. Cells were washed twice with MSM without carbon source and inoculated into fresh MSM at an initial OD_600_ of 0.05 for both monoculture and co-culture conditions. Cultures were incubated at 30 °C with agitation at 300 rpm. Samples (800 µL) were collected hourly over a 10-h period, centrifuged at 13,500 × *g* for 3 min, and the supernatants were analysed by high-performance liquid chromatography (HPLC) to quantify substrate concentrations.

### Construction of synthetic protocatechuate pathway

The synthetic protocatechuate pathway was constructed in pIX-PCA plasmid. Briefly, the *mRFP* reporter gene in the pre-linearized pIX-luxR-mRFP (31) backbone was replaced with four protocatechuate genes: *pcaH*, *pcaG*, *pcaB*, and *pcaC*. These genes were amplified from genomic DNA extracted from ADP1-WT, while the pIX-luxR-mRFP backbone excluding *mRFP* was amplified using primer pair Kan-tlpp-FW/luxR-to-PCA-RV. Assembly was performed using BsaI-based Golden Gate Assembly (GGA), placing the protocatechuate genes under control of the LuxR/P_LuxB_ promoter (32). Ribosome binding sites (RBSs): BCD9, BCD7, and BCD14, previously validated in ADP1 (33), were used to optimize expression. The RBS sequences and their arrangement were selected based on prior evidence of high expression efficiency in ADP1. The resulting plasmid contains a chloramphenicol resistance marker for selection in *E. coli* and a kanamycin resistance marker for use in ADP1. Electrocompetent *E. coli* cells were transformed with assembly mixture; recovered for 1 h at 37 °C in LB medium and plate onto LB agar with chloramphenicol followed by overnight incubation. Colonies were screened by PCR using primers luxR-mRFP-Kan-FW/luxR-mRFP-Kan-RV, and positive clones were verified by Sanger sequencing (Macrogen, The Netherlands). Transformation in ADP1 was performed in ABMS medium and transformants were selected on ABMS agar supplemented with kanamycin and inducer. The integration cassette enabled replacement of the *poxB* locus, a previously characterized neutral site (34), with the protocatechuate gene cluster. Correct chromosomal integration was verified by colony PCR with primer pair poxB-UF/poxB-DR and whole-genome sequencing (WGS). The final strain was designated as ADP1_SYNPCA_.

### Adaptive laboratory evolution (ALE)

To initiate ALE, the ADP1_SYNPCA_ strain was streaked on LB agar supplemented with 1 μM AHL and Km to isolate single colonies. Selected colonies were inoculated into 5 mL minimal salt medium (MSM) containing 0.2% casamino acids, 1 mM PCA, 1 μM AHL, and Km and grown overnight as a seed culture. The overnight cultures were washed twice and resuspended in MSM without carbon source. Cells were then inoculated to an initial OD_600_ 0.1 into 5 mL MSM containing 25 mM PCA as sole carbon source with antibiotic and incubated at 30 °C with shaking at 300 rpm for 2-3 days. Throughout the evolution experiment, 10 μM AHL was used to induce synthetic pathway expression. Cultures were subsequently passaged every 24 h to fresh medium with 25 or 50 mM PCA to a starting OD_600_ of 0.1 for 14 days in total. Glycerol stocks were periodically prepared from the corresponding bacterial cultivations, and individual colonies were isolated by plating the cells on MSM agar containing 25 mM PCA, Km, and 10 µM AHL. The three largest colonies were picked for further characterization. Growth of the evolved clones on PCA was initially assessed in a 96-well plate format in MSM with 25 mM PCA, and the top three performers were validated in 20 mL shake-flask cultures. The cumulative number of generations over the 14-day evolution experiment was calculated using the Eq. (1) (35):

Number of generations = ln(OD_final_/OD_initial_)/ln(2) (1)

### Reverse engineering of ALE-derived mutations

Reverse-engineered strains were constructed by replacing the target locus in the evolved strain using RAMSES as previously described (22), with minor modifications. Correct allele replacement was confirmed by whole-genome nanopore sequencing. The reconstructed strains were subsequently used for phenotypic characterization.

### Analytical procedures

Substrates (benzoate, 4HB, and PCA) were analyzed by HPLC (LC-40D; Shimadzu, Japan) equipped with a photodiode array detector (SPD-M40; Shimadzu, Japan), as previously described (19). Briefly, culture supernatants were filtered through 0.2 µm filters (Agilent Technologies, USA). A Luna C18(2) column (4.6 × 150 mm, 100 Å; Phenomenex, USA) was used and maintained at 40 °C. A mixture of water:methanol:formic acid (80:20:0.16, v/v/v) was used as the mobile phase with the flow rate at 1.0 mL/min. Benzoate, 4-hydroxybenzoate (4HB), and protocatechuate (PCA) were detected at 228, 255, and 259 nm with retention times of 27.25, 7.40, and 4.42 min, respectively. Concentrations were determined from standard calibration curves.

### Whole-genome sequencing

The genomic DNA of ADP1 derived strains and evolved mutants were isolated from LB overnight cultures using a monarch gDNA purification kit (New England Biolabs) or GeneJet genomic DNA purification kit (Thermo Fisher Scientific). The extracted DNA was quantified using a NanoDrop spectrophotometer (Thermo Fisher Scientific) or a Qubit™ 4 Fluorometer with the Qubit™ dsDNA HS Assay Kit (Thermo Fisher Scientific). Samples were sequenced using nanopore technology. Libraries were prepared with the Q20+ Duplex-Enabled Rapid Sequencing Kit (Oxford Nanopore Technologies, UK) according to the manufacturer’s instructions. Sequencing was carried out on a MinION device using a Flow Cell R10 (FLO-MIN114) (Oxford Nanopore Technologies), operated with MinKNOW software (Oxford Nanopore Technologies). The bacterial genome sequences were analyzed with *breseq* (36) to predict point mutations, large deletions, and other types of structural variation relative to the ADP1 genome and theoretical genome sequence of ADP1_SYNPCA_ parent strain.

## RESULTS

### Construction of *A. baylyi* ADP1 deletion strains

ADP1 harbours a highly branched and complex aromatic degradation network, composed of multiple peripheral pathways that funnel diverse substrates into the β-ketoadipate pathway (KAP), a central catabolic node for aromatic compound assimilation (37–39). The KAP comprises three main branches: protocatechuate (PCA), catechol (CAT), and dicarboxylic acids (DCA) route. The protocatechuate and catechol branches converge at β-ketoadipate enol-lactone, which is subsequently converted to tricarboxylic acid (TCA) cycle intermediates, namely acetyl-CoA and succinyl-CoA (26). In parallel, the dicarboxylic acids branch channels carboxylic acids to β-ketoadipyl-CoA, merging downstream into the main KAP route. This modular yet intertwined architecture underlies the ADP1’s broad catabolic versatility but simultaneously complicates metabolic rewiring for lignin-related aromatic compound utilization and valorization. In particular, overlapping substrate specificities, shared transport systems, and cross-regulatory networks blur functional boundaries between branches, limiting the predictability of pathway refactoring and flux redistribution.

To simplify this complexity and enable pathway-level dissection, we employed a dual-chassis strategy using ADP1 wild-type (ADP1-WT) and its stable genome-derivatives ISx and ISx_csrA_. ISx_csrA_ is derived from ISx. It restores a truncation of the global carbon regulator *csrA* that is present in ADP1-WT (40). ISx, a genetically stable and transposon-free chassis was used together with ISx_csrA_ to assess the contribution of this global regulator to growth-related phenotypes. These strains were selected as complementary chassis: ADP1-WT serves as the standard reference with native regulatory complexity and adaptability, while ISx/ISx_csrA_ provides a stable genome variant optimized for genetic stability. This sequential framework was designed to generate: i) a minimal aromatic degradation network through targeted removal of redundant and competing routes, ii) evolved ADP1-WT variants capturing adaptive mutations that restore fitness and metabolic balance following streamlining, and iii) a stabilized platform that incorporates optimized pathway architecture and adaptive features within a genome-stabilized background suited for further refactoring and more efficient and comprehensive lignin-derived substrate utilization.

Building on these concepts, we first created branch specific deletion strains in which either the entire catechol, protocatechuate, or dicarboxylic acid branch — including the respective upper funneling pathway — was removed. This strategy allowed us to evaluate each branch independently, without interference from parallel pathways or overlapping regulatory systems. Markerless deletions were generated by Golden Transformation to construct deletion strains in ADP1, ISx, and ISx_csrA_. The catechol branch deletion strain (ΔCAT) was constructed by removing anthranilate (*antA, antB, antC)* and *sal-are-ben-cat* gene set (ACIAD1424 to ACIAD1451), resulting in a ∼31.9-kb chromosomal deletion. Similarly, the protocatechuate branch deletion strain (ΔPCA) was generated by removing genes associated with the metabolism of aromatic substrates funneled into the protocatechuate branch. This cluster includes the *van* (ACIAD0978 to ACIAD0983) and *pca-qui-pob-hca* genes (ACIAD1702 to ACIAD1728). Lastly, although adipate is not processed via the canonical aromatic branches, its catabolism converges with the central trunk of the β-ketoadipate pathway (41). Therefore, for the dicarboxylic acid branch, we deleted the *mucK* muconate transporter (42) and *dca* cluster region (ACIAD1681 to ACIAD1699) to eliminate their contributions to aromatic catabolism. The strain carrying a deletion in the dicarboxylic acid branch was designated as ΔDCA. Finally, we combined the ΔPCA, ΔCAT, and ΔDCA, deletions to entirely abolish the β-ketoadipate pathway, in a strain designated ΔKAP, which has a total of ∼87.1 kb deleted from the chromosome. The deletion strains were verified by colony PCR and whole-genome sequencing. The resulting strains provided a minimal yet functional framework for characterizing aromatic substrate utilization and served as the basis for subsequent synthetic operon integration and adaptive laboratory evolution. Fig. 1 summarizes the genomic modifications performed in each strain and maps the deleted regions to their chromosomal coordinates in ADP1.

**Figure 1.**
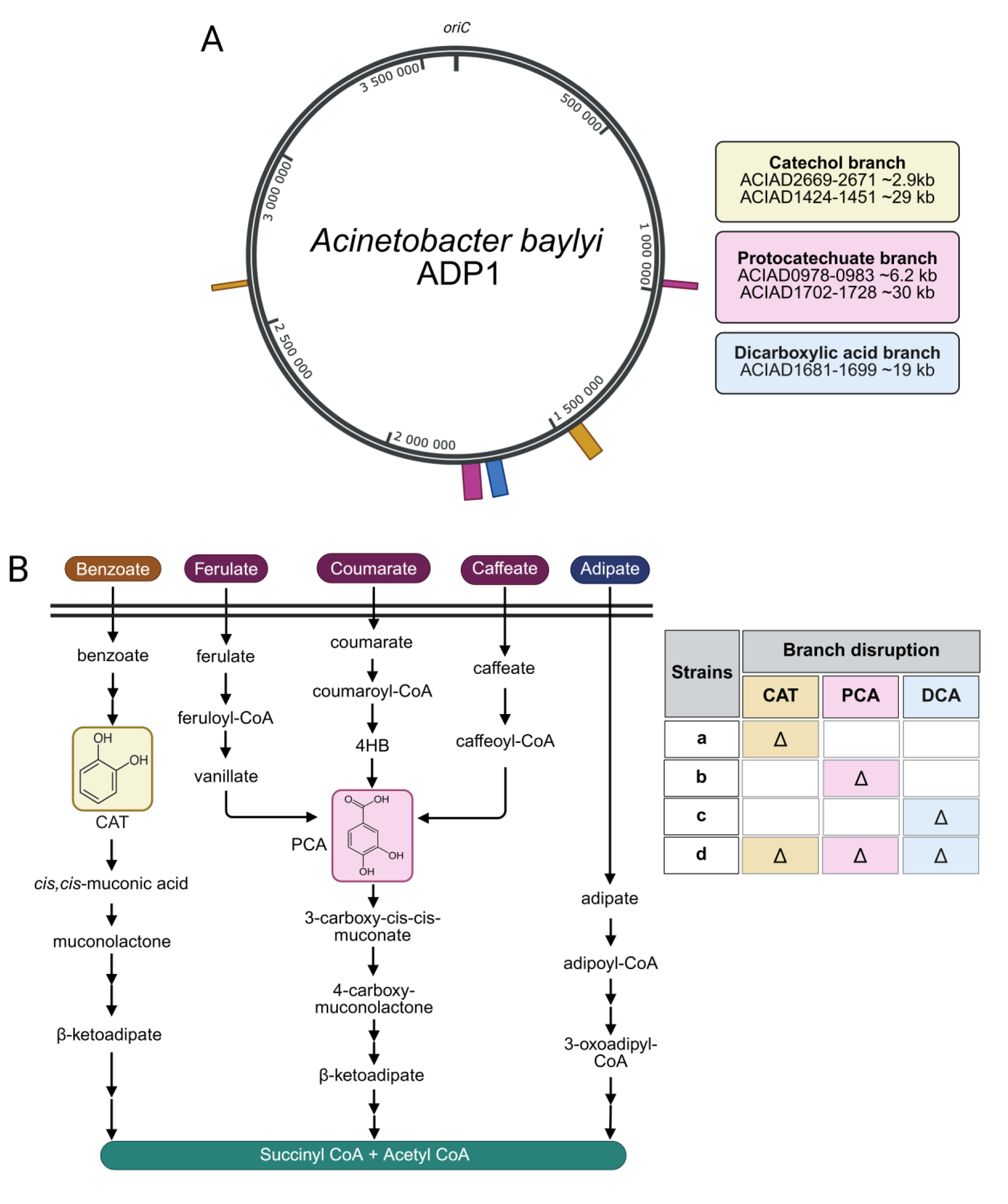
Construction of deletion strains in ADP1, ISx, and ISx_csrA_. A) The chromosomal locations and sizes of the gene clusters (catechol, protocatechuate, and dicarboxylic acid branch) responsible for aromatic compound catabolism in ADP1. B) Simplified schematics of the aromatic degradation pathways in ADP1. Δ; deletion, CAT: catechol; PCA: protocatechuate; DCA: dicarboxylic acids; KAP: β-ketoadipate pathway. Example substrates are color-coded according to their corresponding pathway branches: catechol (CAT, yellow), protocatechuate (PCA, pink), and dicarboxylic acid (DCA, blue).

### Characterization of ADP1 deletion strains

We next assessed the catabolic capabilities of deletion strains to utilize aromatic compounds by evaluating growth on six lignin-related model compounds as sole carbon sources across a range of concentrations. ADP1-WT, ISx, ISx_csrA_, and the deletion strains (ΔCAT, ΔPCA, ΔDCA, and ΔKAP) were initially cultured in MSM supplemented with 0.2% casamino acids and 1 mM of the target compound to allow initial adaptation. Cells were then pelleted, washed, and transferred to MSM medium containing only the test compound. Growth kinetics and lag times were measured following the transfer, providing insights into each strain’s ability to convert the compounds into biomass and energy, as well as their tolerance to these lignin-derived substrates.

The strains with individual branches deleted were not able to grow on the corresponding substrates catabolized in these pathways: in the 48 h growth assays using the defined substrates, the catechol-deletion strain (ΔCAT) was unable to grow on benzoate, while the protocatechuate-deletion strain (ΔPCA) was unable to utilize 4-hydroxybenzoate (4HB), *p*-coumarate, *trans*-ferulate, and vanillate. Similarly, the dicarboxylic acid-deletion strain (ΔDCA) showed no growth on adipate. As expected, the triple-deletion strain (ΔKAP), lacking all three pathways, did not grow on any of the tested compounds, confirming that each pathway is essential for its corresponding substrate (Fig. 2). Furthermore, quantitative analysis revealed that 25 mM benzoate inhibited growth across all strains, indicating substrate toxicity at high concentrations. In contrast, the strains in which the protocatechuate branch was intact were able to grow in the presence of 4HB and vanillate up to 25 mM without detectable inhibition (μ_max_=0.45-0.51 h^-1^; lag phase 5-8.5 h). The *p*-coumarate and *trans*-ferulate, however, impaired growth at this level, with *trans*-ferulate causing a pronounced delay in growth already at 10 mM in both wild-type and deletion strains, resulting in an extended lag phase of approximately ∼20 h compared to *p*-coumarate (Supplementary Fig. S3-S5). Notably, the ISx strain exhibited a markedly long lag phase (∼20 h) on 25 mM vanillate, compared to 10 h observed for ADP1-WT and ISx_csrA_. Overall, deletion of a single branch (protocatechuate or catechol or dicarboxylic acids), did not affect growth on the substrate metabolized by the remaining branches, indicating that each route operates independently under the tested conditions.

**Figure 2.**
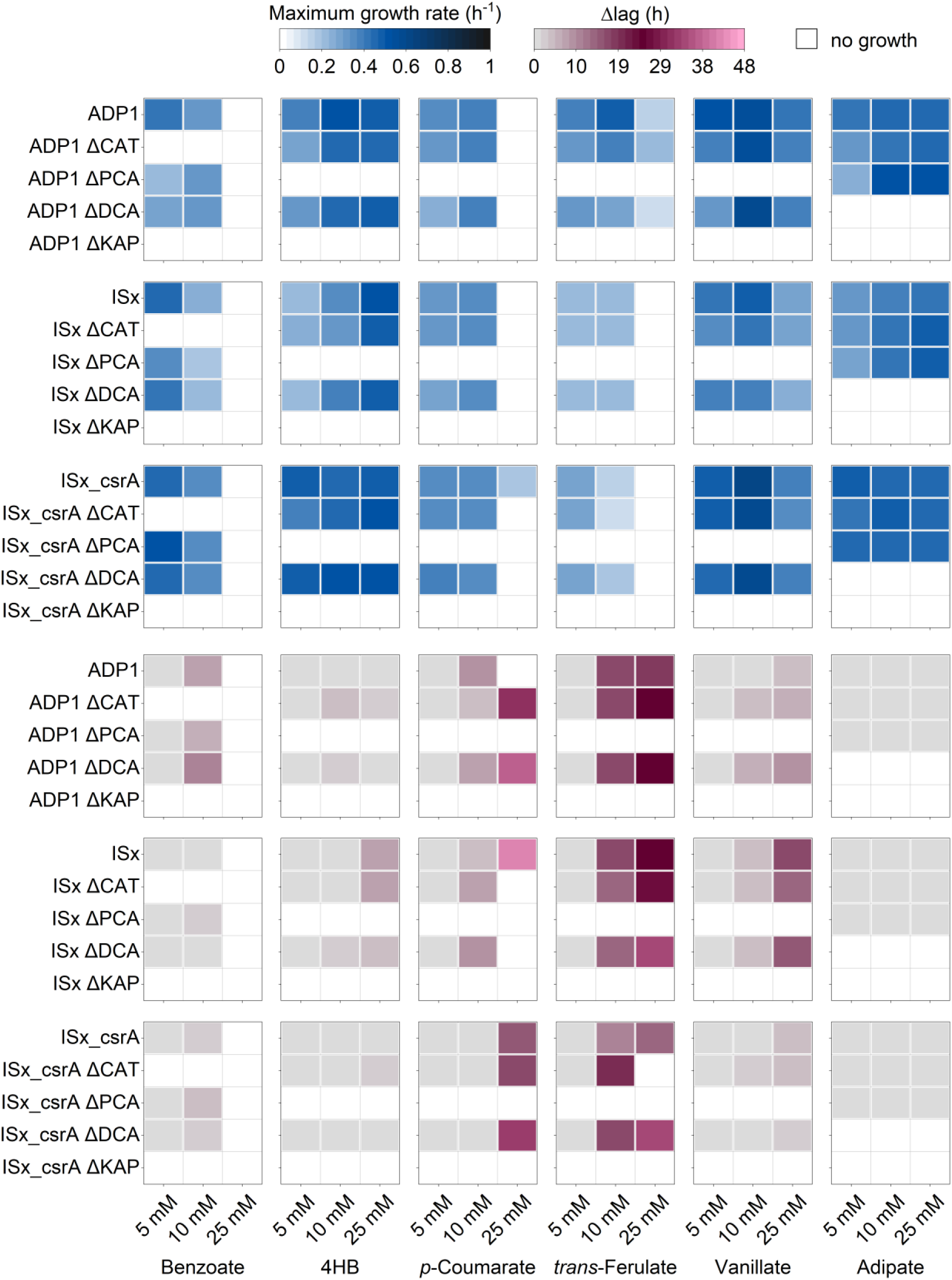
Growth profiles of ADP1-WT, ISx, ISx_csrA_ and the constructed deletion strains (ΔCAT, ΔPCA, ΔDCA, and ΔKAP). Bacterial growth in 96-well plates containing 200 µL of growth medium was monitored on Spark multimode microplate reader (Tecan, Switzerland) for 48 hours. Strains were grown in MSM supplemented with different concentration of benzoate, 4HB, *p*-coumarate, *trans*-ferulate, vanillate, or adipate as sole carbon sources ranging from 5 mM to 25 mM. Growth curves and detailed values are provided in Supplementary Figure S3-S5 and Supplementary Data 1, respectively. The data represents the mean of the calculated growth rate and Δlag times determined from three biological replicates.

### Utilization of mixed aromatic substrates

In order to determine whether substrate utilization by the deletion strains (ΔCAT or ΔPCA) is affected by the presence of another aromatic compound, we tested the strains in mixed-substrate conditions under two setups: 1) 10 mM 4HB with different concentrations of benzoate, and 2) 5 mM benzoate with different concentrations of 4HB. In the first experiment, all ΔCAT strains exhibited growth inhibition at higher benzoate concentrations, indicating a dose-dependent toxic effect (Fig. 3A). To determine whether this effect was carbon-source dependent, we next evaluated growth on gluconate, which ADP1 readily metabolizes. Growth on 50 mM gluconate was similarly reduced in the presence of elevated benzoate, supporting a general toxic effect of benzoate (Supplementary Fig. S6). In contrast, when 5 mM benzoate was combined with different concentrations of 4HB, no toxic effects were observed on ΔPCA strains (Fig. 3B). Notably, benzoate appeared to be more toxic to ISx strains than to ADP1-WT. Furthermore, in the ISx_csrA_ strains, growth remained inhibited in the presence of 3 mM benzoate, but the extent of inhibition was slightly reduced compared to the ISx, suggesting a modest reduction in benzoate sensitivity.

**Figure 3.**
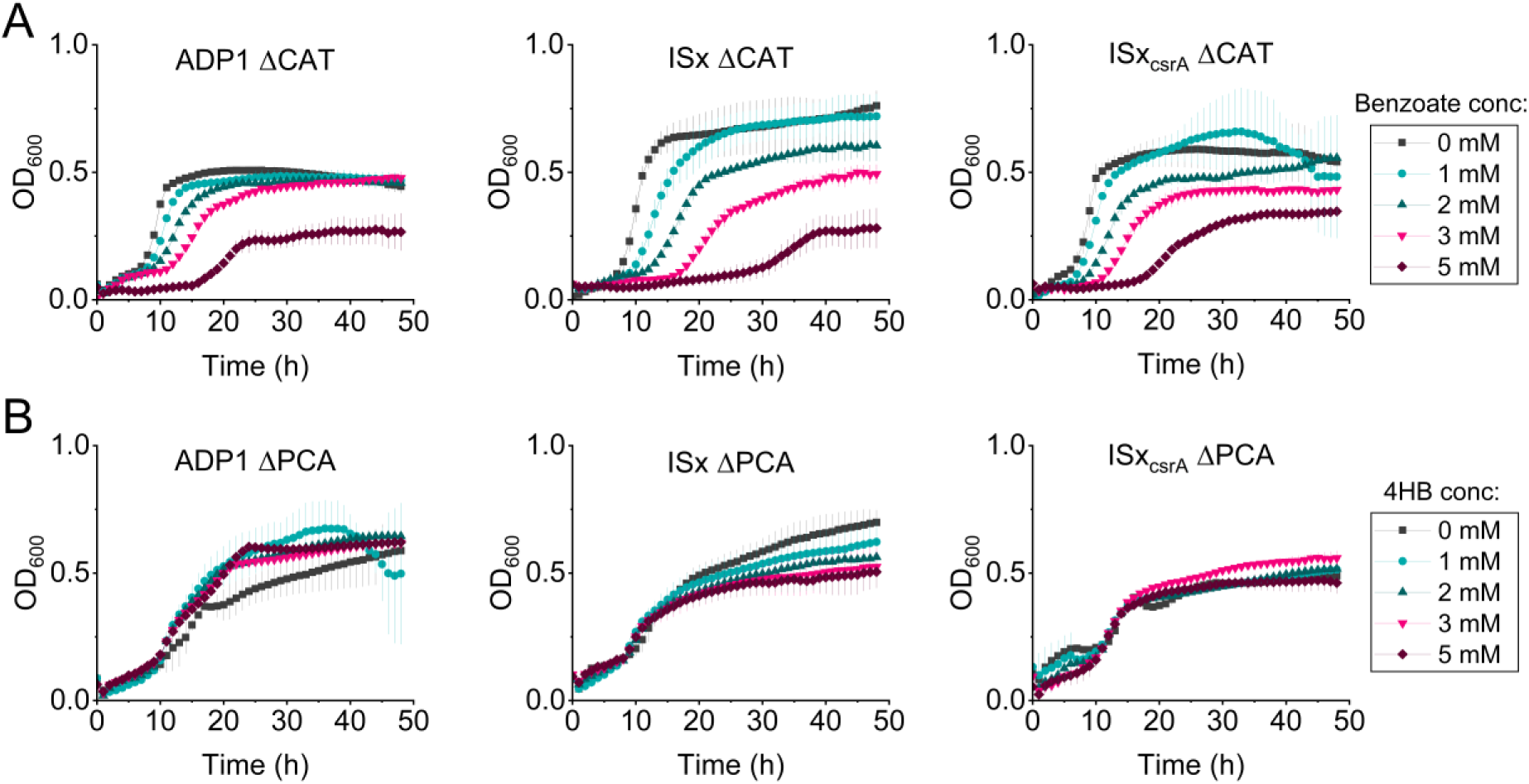
Evaluation of benzoate and 4HB toxicity in ADP1, ISx, and ISx_csrA_ deletion strains. (A) Growth of the catechol branch deletion strains (ΔCAT) in ADP1, ISx, and ISx_csrA_. Cultures were grown in minimal salts medium supplemented with 10 mM 4HB and 0, 1, 2, 3, or 5 mM benzoate. Growth curves showing reduced growth at increasing benzoate concentrations. B) Growth of the protocatechuate branch deletion strains (ΔPCA) in ADP1, ISx, and ISx_csrA_. Cultures were grown in minimal salts medium supplemented with 5 mM benzoate and 0, 1, 2, 3, or 5 mM 4HB. Cell growth was monitored by measuring optical density at 600 nm (OD_600_) every hour for 48 hours using a microplate reader. Data represents the mean ± standard deviation of three biological replicates (n = 3).

We next evaluated the performance of the deletion strains in co-culture, focusing on the ADP1ΔCAT – ADP1ΔPCA pair, which exhibited higher tolerance to benzoate under mixed-substrate conditions. Each deletion strain selectively consumed the substrate associated with its intact pathway, while the utilization of the target substrate was unaffected by removal of the other pathways (Supplementary Fig. S7). Based on this behaviour, we hypothesized that pairing these strains could enable simultaneous substrate consumption through functional partitioning between members. Although cooperative degradation has previously been demonstrated in engineered *A. baylyi* consortia (43), our goal was to investigate how these deletion strain pairs perform under mixed-substrate conditions and whether they can be leveraged to establish a stable, modular consortium (Fig. 4A).

**Figure 4.**
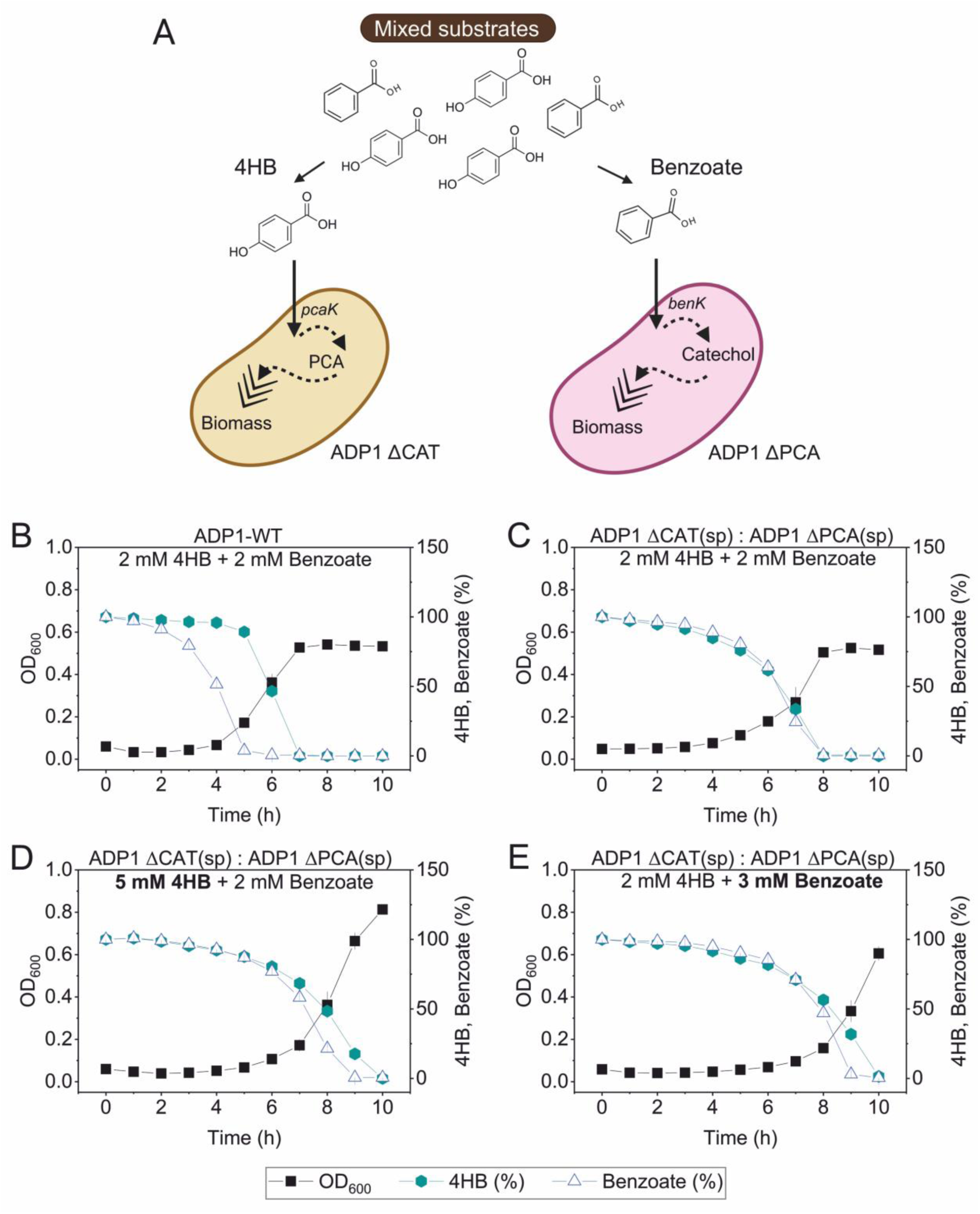
Co-culture strategy for mixed substrates utilization. A) Schematic representation of utilization of mixed substrates by two ADP1 deletion strains. B) Monoculture of ADP1-WT in mixed substrate 2 mM 4HB and 2 mM benzoate. Growth curves showing preferential substrate consumption of benzoate followed by 4HB. C) D) E) Represents the coculture of ADP1 ΔCAT(sp) and ADP1 ΔPCA(sp) strains at a 1:1 inoculation ratio in minimal salts medium (MSM) supplemented with varying concentrations of 4HB and benzoate. Growth conditions were as follows: (C) 2 mM 4HB and 2 mM benzoate, (D) 2 mM 4HB and 3 mM benzoate, and (E) 5 mM 4HB and 2 mM benzoate. Optical density at 600 nm (OD_600_) and substrate consumption (%) were measured every hour for 10 hours. Substrate concentrations were quantified by high-performance liquid chromatography (HPLC). Data represents the mean ± standard deviation of three biological replicates (n = 3). Strains carrying CRISPR spacers are indicated by “(sp)”.

ADP1 exhibits high natural transformation efficiency, which can lead to spontaneous reacquisition of deleted DNA regions during co-culture. To prevent this and maintain genetic stability, we implemented a CRISPR-Lock system that targets deleted loci for CRISPR-mediated interference (25). A CRISPR-ready module was first introduced into ADP1 ΔCAT and ADP1 ΔPCA, replacing the native CRISPR array with a *tdk–kanR* cassette (25), enabling subsequent integration of synthetic spacers. Two spacers were designed: i) a cat-spacer targeting deleted region in the CAT pathway, ii) a pca-spacer targeting deleted region in the protocatechuate pathway. The CRISPR-ready variants were then rescued with either a cat or pca spacer, each targeting 32 bp of the deleted region, thereby preventing reacquisition of the corresponding genes from the partner strain. This system ensured that division of labour remained stable by blocking genetic complementation through natural transformation (25). The correct genomic integration was then confirmed by colony PCR and kanamycin sensitivity testing. The resulting strains were designated as ADP1 ΔCAT(sp) and ADP1 ΔPCA(sp), carrying cat and pca-spacer, respectively. With this safeguard in place, we next evaluated the strains in co-culture. The strains containing spacers were pre-adapted in minimal medium with 0.2% casamino acids and a mixture of aromatics (1 mM of each), then transferred to carbon-free medium prior to inoculation. Mixed-substrate experiments were performed with different combinations of benzoate and 4HB, and growth and substrate consumption were monitored over time by OD_600_ and HPLC measurements.

As we expected, the monoculture ADP1-WT strain consumed mixed substrate in a sequential manner where benzoate was consumed first, followed by 4HB consumption (Fig. 4B). In contrast, the co-culture ADP1 ΔCAT(sp) : ADP1 ΔPCA(sp) (1:1) depleted both substrates simultaneously, demonstrating parallel utilization without detectable regulatory interference. Furthermore, simultaneous consumption of benzoate and 4HB by the co-culture was consistently demonstrated across all tested conditions (Fig. 4C-4E).

After confirming that the CRISPR-Lock single-deletion strains selectively consumed their target substrates without inhibition, we next tested whether this design could be extended. For this, we constructed double deletion ADP1 variants lacking both the CAT and DCA branches (ΔCAT ΔDCA) or the PCA and DCA branches (ΔPCA ΔDCA), each equipped with the corresponding CRISPR-Lock spacer (designated ADP1 ΔCAT ΔDCA(sp) and ADP1 ΔPCA ΔDCA(sp), respectively). These double deletion strains were tested in co-culture to evaluate their ability to consume aromatic substrates and to determine whether the presence of another substrate introduced any inhibitory effects. Similarly, co-culture experiments demonstrated concurrent utilization of mixed substrates (Fig. 5A). However, in the presence of 2 mM benzoate and 2 mM 4HB, a modest delay was observed in benzoate utilization. Increasing either benzoate or 4HB concentrations did not prevent simultaneous utilization of mixed substrates. These results demonstrate that co-culturing functionally specialized strains enable parallel degradation of aromatic substrates that are otherwise consumed sequentially in the wild-type strain.

**Figure 5.**
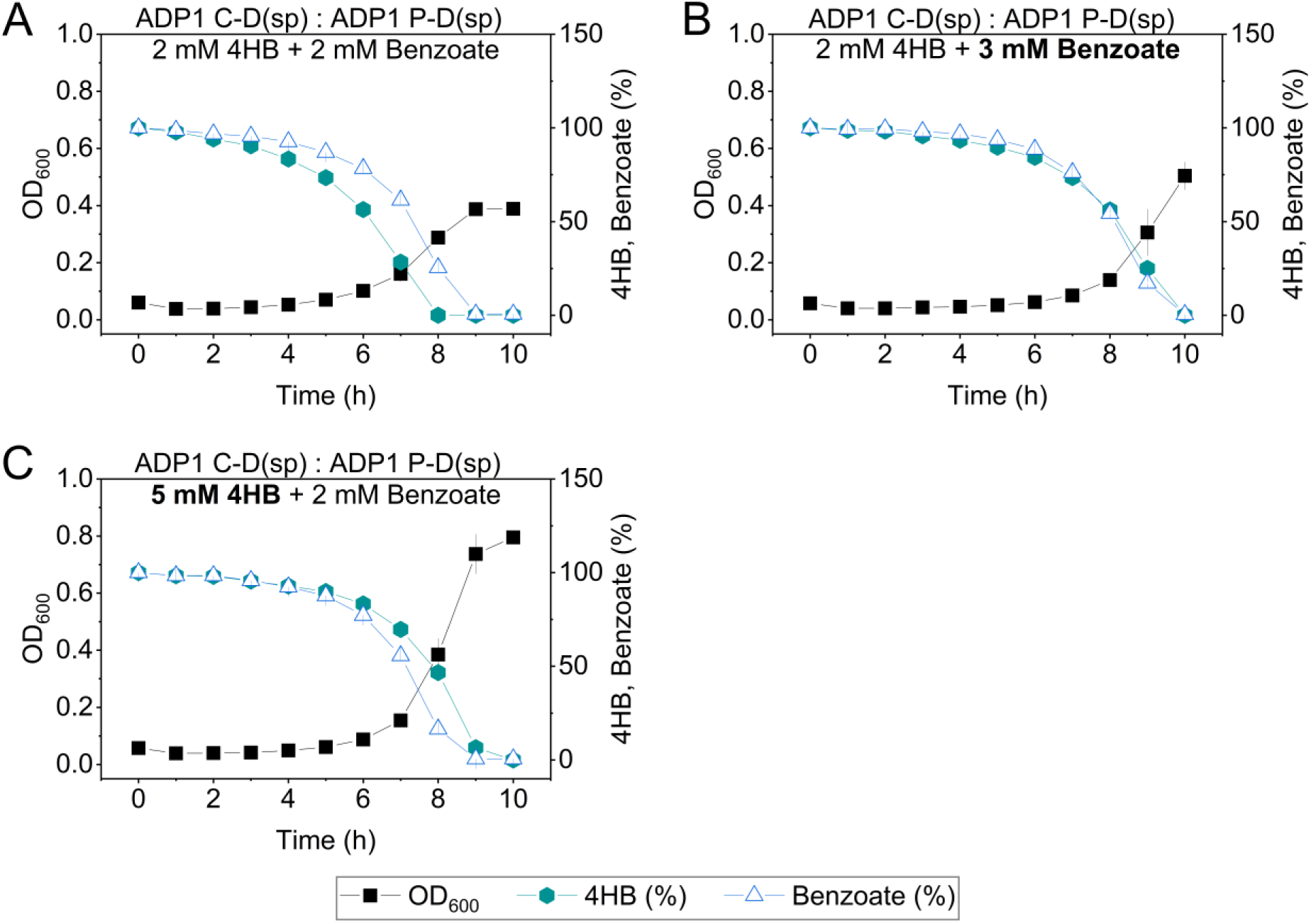
Coculture of ADP1 ΔCAT ΔDCA(sp) and ADP1 ΔPCA ΔDCA(sp) strains at a 1:1 inoculation ratio in minimal salts medium (MSM) supplemented with varying concentrations of 4HB and benzoate. Growth conditions were as follows: (A) 2 mM 4HB and 2 mM benzoate, (B) 2 mM 4HB and 3 mM benzoate, and (C) 5 mM 4HB and 2 mM benzoate. Optical density at 600 nm (OD_600_) and substrate consumption (%) were measured every hour for 10 hours. Substrate concentrations were quantified by high-performance liquid chromatography (HPLC). Data represent the mean ± standard deviation of three biological replicates (n = 3). For clarity, strain names are abbreviated in figure labels (ADP1 C-D, ADP1 ΔCAT ΔDCA; ADP1 P-D, ADP1 ΔPCA ΔDCA), and strains carrying CRISPR spacers are indicated by “(sp)”.

### Synthetic pathway for integrating protocatechuate utilization into the catechol branch

To streamline the catabolism of the key intermediates, the protocatechuate branch, previously operating in parallel, was merged into catechol branch (Fig. 6A). We reconstructed this route using a minimal set of essential genes for protocatechuate utilization under inducible control. This design aimed to accomplish three goals: 1) to merge the catechol and protocatechuate entry points into a unified catabolic module, 2) to remove protocatechuate native regulatory constraints and redundant genes in KAP, and 3) to test whether the rewired configuration could support efficient growth on protocatechuate through the catechol pathway. This restructuring effectively redirected the metabolic flux through a single channel, allowing us to assess the performance and stability of minimal pathway architecture.

**Figure 6.**
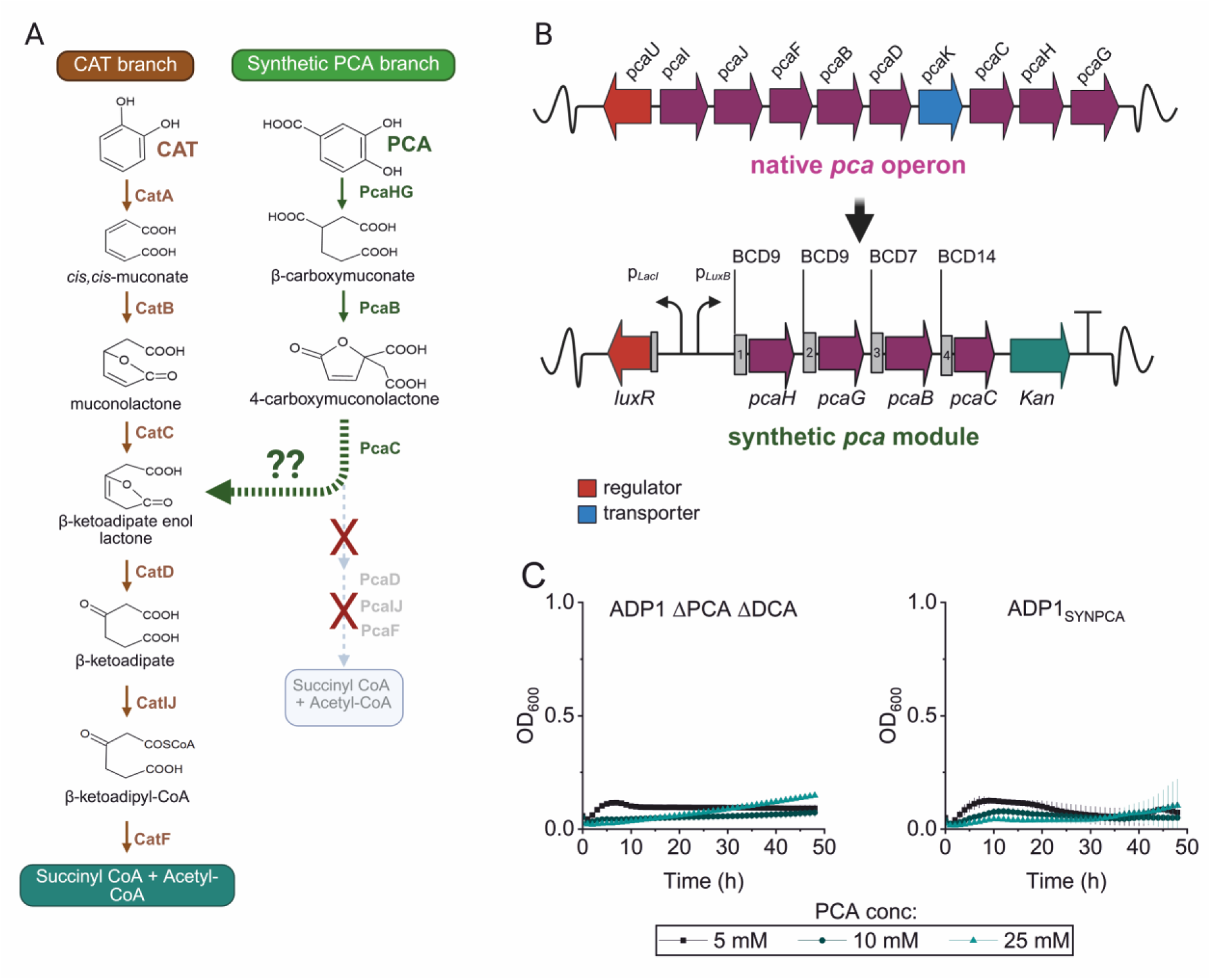
Integration of synthetic protocatechuate pathway into the catechol branch. A) Schematic representation of the reconstructed aromatic degradation pathway, B) Comparison of the native *pca* (∼9.1 kb) vs synthetic *pca* genes under LuxR/P_LuxB_ promoter (∼3.1 kb) in ADP1 ΔPCA ΔDCA genome. C) Growth curve of ADP1 ΔPCA ΔDCA and ADP1_SYNPCA_ in MSM with different concentration of PCA (5, 10, and 25 mM). ADP1_SYNPCA_ cultures were induced with 10 μM of AHL. Data points represent means ± standard deviations of biological replicates (n = 3). CAT: catechol, PCA: protocatechuate.

Next, to identify a suitable genomic context for stable and consistent expression, we tested two chromosomal loci: location I (*poxB*, ACIAD3381), a well-characterized neutral site (17, 44), and location II (*pca-qui-pob-hca* gene cluster, ACIAD0983-ACIAD1702). The suitability of each site for gene expression was assessed by fluorescent reporter mRFP (45) under the regulation of LuxR/P_luxB_ (32). At location II, two orientations of the construct were tested; however, only one supported growth on LB or minimal salts medium (MSM) supplemented with antibiotics, suggesting that the alternative configuration likely led to incorrect orientation of the integrated cassette. Fluorescence levels were subsequently quantified using a microplate reader. Location I exhibited higher mRFP expression compared to location II (Supplementary Fig. S8) and showed a LuxR-dependent response to the inducer, indicative of tighter transcriptional control and greater regulatory responsiveness. Based on this expression profile, location I was selected as the integration site.

A synthetic protocatechuate pathway was designed by introducing only the essential genes required for its utilization (Fig. 6A). These genes consist of *pcaG*, *pcaH*, *pcaB*, and *pcaC* which encode key enzymes in the protocatechuate branch of the β-ketoadipate pathway. The construct excluded genes encoding for a transporter (*pcaK*), regulator (*pcaU*), and genes encoding for functions already present in the catechol branch (*pcaDIJF*). Furthermore, in the native configuration, the genes are arranged as *pcaBCHG* (Fig. 6B), whereas our synthetic operon was restructured as *pcaHGBC*, thereby prioritizing the first step of PCA conversion. All genes were PCR-amplified from ADP1-WT genome, placed under the control of an inducible lux promoter (LuxR/P_luxB_) to allow tunable expression (32) and equipped with ribosome binding sites (RBSs) BCD9, BCD14, and BCD7 (33). The RBS sequences and their arrangement were chosen based on prior evidence of high expression efficiency in ADP1 (33). The synthetic protocatechuate module was flanked by sequences homologous to the genome integration site replacing ACIAD3381. The construct was assembled in a plasmid containing Km and Cm resistance markers for selection in ADP1 and *E. coli* XL-1, respectively, and propagated in *E. coli* for sequence verification. The verified plasmid that is non-replicative in ADP1 was then introduced into the genome of ADP1 ΔPCA ΔDCA by natural transformation, and transformants were selected on Km. Candidate colonies were screened by colony PCR and confirmed by Sanger sequencing, followed by whole-genome nanopore sequencing to verify correct chromosomal integration. The strain was designated as ADP1_SYNPCA_.

To evaluate the functionality of the engineered strain, we tested its growth in MSM supplemented with varying concentrations of PCA as a sole carbon source. No detectable growth was observed under any tested condition (Figure 6C). Further supplementation with casamino acids at 1% both in the presence and absence of PCA, resulted in similar final optical densities (OD_600_), indicating that PCA was not consumed under these conditions (Supplementary Fig. S9). These results indicate that the engineered strain was unable to utilize PCA, possibly due to incomplete restoration of essential genes required for protocatechuate uptake or catabolism.

### Adaptive Laboratory Evolution of ADP1_SYNPCA_

Given the insufficient growth of ADP1_SYNPCA_ on PCA, we applied ALE under selective pressure in MSM with PCA as the sole carbon source (Fig. 7A). More than ten independent colonies were initially screened for growth under selective conditions, and two colonies that sustained growth were subsequently propagated as independent evolutionary lines. Cultures were first incubated for 72 hours to permit acclimation, followed by daily passage for 14 consecutive days, corresponding to approximately 60 generations (Fig. 7B, Supplementary Table S7). Evolved populations from two independent lineages were plated on MSM agar supplemented with 25 mM PCA at three key evolutionary stages: early (E, day 3), mid (M, day 7), and late (L, day 14). To identify improved variants, a larger number of colonies (n = 13 per lineage) were screened at the late stage, where adaptive mutations are more likely to have accumulated. From these, the top three colonies from L stages were selected based on growth performance (Supplementary Fig. S10), while three representative isolates were also chosen from each of the early and mid-stages, yielding a total of nine candidates for further characterization. To evaluate whether growth performance differed across the evolutionary stages, all isolates were cultivated in MSM with 25 mM PCA as the sole carbon source, and growth was monitored every hour for 48 hours using a microplate reader.

**Figure 7.**
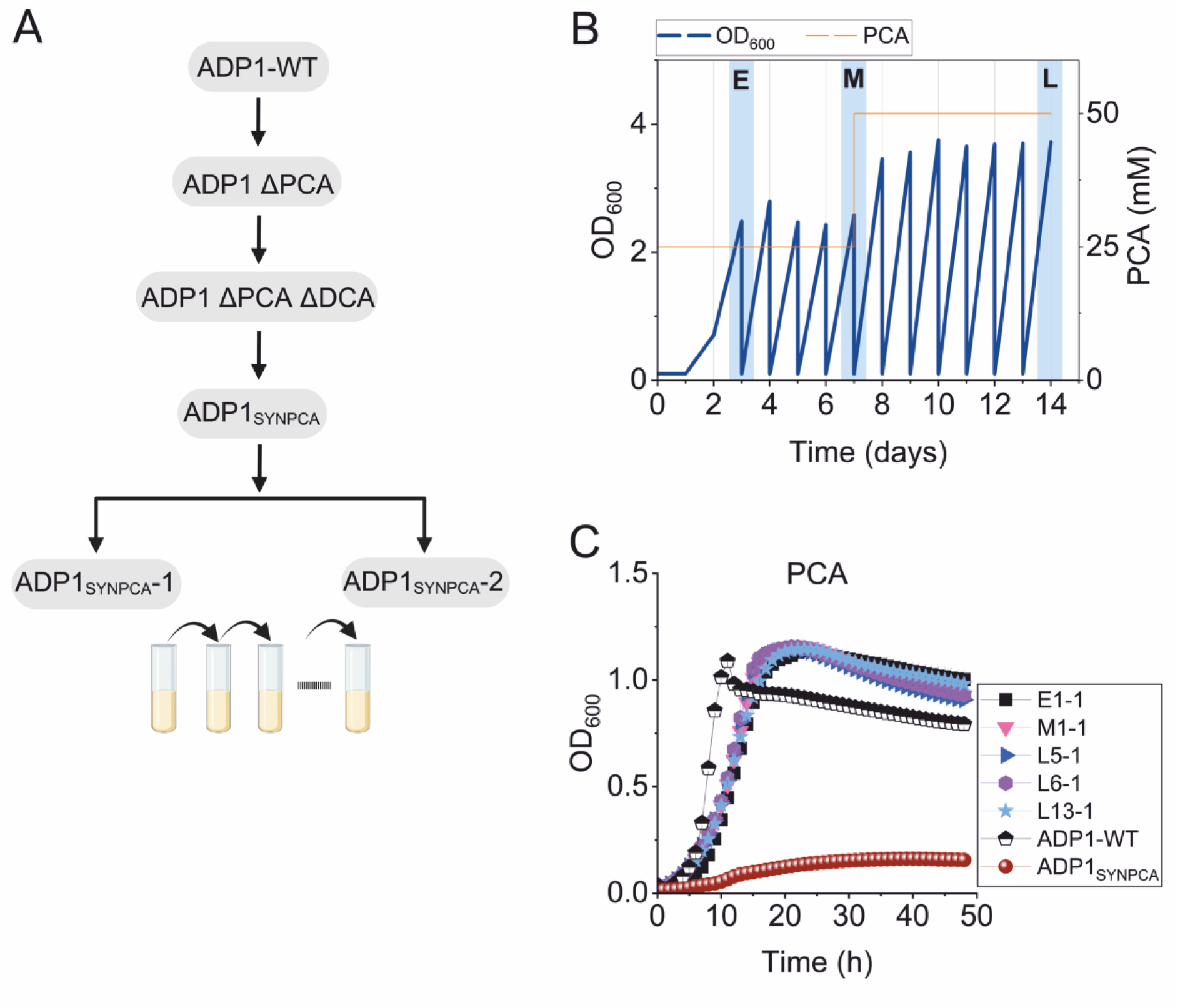
Adaptive laboratory evolution (ALE) of ADP1_SYNPCA_ on PCA. A) Dendogram of ADP1 strains for ALE. B) OD_600_ during ALE on PCA for 14 days. Strains in early (E), mid (M), and late (L) evolution stages were shown in blue shades. PCA concentration was increased from 25 mM to 50 mM and the glycerol stock was prepared every day after 72h. The data are presented as the average of two evolutionary lines. B) Growth profiles of evolved strains from ADP1_SYNPCA_-1 (E, M, and L) in MSM with 25 mM PCA with Km and 10 μM of AHL. Data points represent the mean ± standard deviation from *n* = 3 biological replicates. Details of the growth profiles of the evolved strains are shown in Supplementary Fig. S11.

All evolved isolates exhibited growth on PCA as a sole carbon source (Fig. 7C, Supplementary Fig. S11). The growth profiles revealed no substantial differences between early (E) and mid-stage (M) isolates, whereas late-stage (L) strains exhibited a modestly shorter lag on PCA. This indicates that mutations enabling PCA utilization likely arose already during the early phase of evolution, with only minor refinement in later stages. All evolved strains displayed lag phases of ∼4–6 h, with a small reduction in the L strains (4 h compared to 5–6 h in earlier isolates). Notably, while early isolates exhibited a longer lag phase, they had a slightly higher μ_max (0.32-0.38 h⁻¹) compared to mid/late isolates (0.22-0.25 h⁻¹). This elevated μ_max reflects only a short-lived advantage during exponential growth, without affecting average growth rate or final biomass yield. In agreement, μ_average was somewhat higher in early isolates (0.25–0.30 h⁻¹) than in mid/late isolates (0.20–0.23 h⁻¹), but all values remained below the wild type. OD_max values were comparable across lineages (1.1 versus 1.0 for ADP1-WT; Supplementary Data 2).

To assess whether mutations acquired during evolution had broader effects on carbon metabolism, we tested the growth of evolved strains on alternative carbon sources: benzoate, succinate, and glucose. These substrates were selected to probe distinct catabolic pathways and to identify potential trade-offs or cross-pathway effects associated with adaptation. Growth phenotypes varied across substrates, with late evolved strains exhibiting profiles comparable to the ADP1-WT and parental strain ADP1_SYNPCA_ under each condition (Fig. 8). Similar behavior was observed across independent evolutionary lineages (Supplementary Fig. 12). On benzoate, both μ_max (0.11–0.21 vs 0.25 h⁻¹) and μ_average (0.10–0.16 vs 0.22 h⁻¹) were clearly reduced relative to ADP1-WT, although OD_max was similar (∼0.7-0.8). On succinate, μ_max values of L strains overlapped with ADP1-WT (0.34–0.44 vs 0.47 h⁻¹), but μ_average was consistently lower (0.22–0.32 vs 0.42 h⁻¹), while OD_max remained at similar level (0.9–1.0 vs 1.0). In contrast, on glucose no significant differences were observed between evolved isolates and ADP1 across μ_max, μ_average, or OD_max, indicating that the mutations did not substantially affect other catabolic functions. We also tested the L strains on MSM supplemented with 25 mM PCA as the sole carbon source under varying concentrations of AHL inducer to assess whether growth was induction dependent. The results revealed a dose-dependent response; OD_600_ were comparable across inducer concentrations ranging from 1 to 100 μM (Supplementary Fig. S13). In contrast, growth was slower in the absence of inducer (0 μM), indicating that induction is required for optimal performance. This observation is consistent with earlier *mRFP* reporter assays, which showed saturated promoter activity at concentrations above 10 μM AHL, suggesting that full pathway induction had already been achieved at low inducer levels.

**Figure 8.**
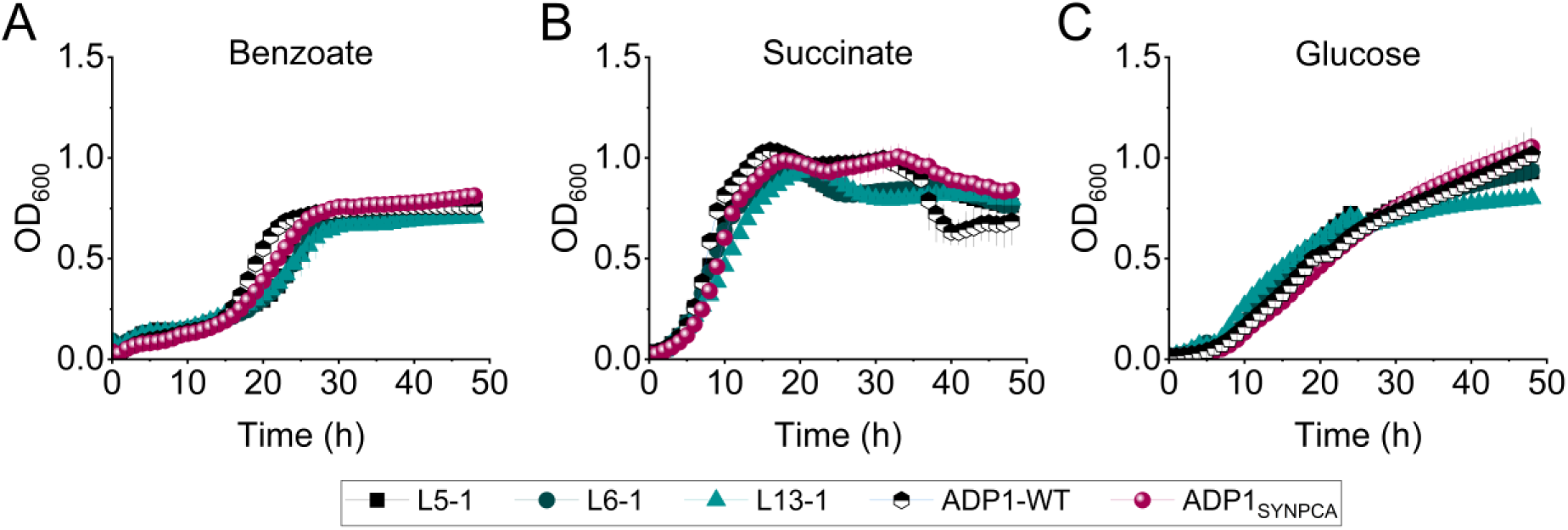
Growth profiles of the late evolved strains (L) from evolutionary line 1 compared to ADP1-WT and parental strain ADP1_SYNPCA_ in MSM supplemented with different carbon sources: A) 10 mM benzoate, B) 50 mM succinate, C) 50 mM glucose. Data points represent the mean ± standard deviation from *n* = 3 biological replicates. An independent evolutionary lineage (line 2) showed comparable behaviour (Supplementary Fig. 12). Underlying data and calculated growth parameters are provided in Supplementary Data 3.

To further assess strain performance and monitor PCA utilization, we performed flask cultivations using 50 mM PCA as the sole carbon source. Cultures were incubated for 30 hours, and samples were collected every 3 hours to monitor growth and substrate consumption. The OD_600_ and substrate concentrations were measured at each time point using spectrophotometer and HPLC, respectively. All evolved strains successfully utilized the compound, reaching maximal cell densities between 15 and 18 hours, with peak OD_600_ values around 5. L13-1 and the ADP1-WT strains peaked at 18 hours, while the remaining strains reached maximal OD_600_ slightly earlier at 15 hours. These results confirm that the evolved strains retained efficient substrate utilization comparable to the wild-type strain under elevated substrate conditions (Supplementary Fig. S14).

### Evaluation and reverse engineering of the ALE-derived mutations

To identify mutations potentially contributing to PCA utilization, we sequenced the genomes of ten evolved isolates representing three time points from two evolutionary lines (Supplementary Table S8). Nine out of ten isolates carried an identical synonymous mutation in muconate cycloisomerase I *catB* (R257R; CGA→AGA) (Fig. 9A), which is responsible for converting *cis,cis*-muconate to muconolactone in the catechol pathway. To assess the functional contribution of this mutation, we replaced the *catB* locus in the evolved strain L5-1 with either the wild-type or the evolved sequence, generating L5-1-catB^WT^ and L5-1-catB^R257R^, respectively. Five independent colonies of each strain were selected and evaluated for growth in MSM supplemented with 25 mM PCA, Km, and 10 µM AHL. In microplate reader assays, L5-1-catB^WT^ failed to grow on PCA as the sole carbon source, whereas both the evolved strains L5-1 and L5-1-catB^R257R^ retained the ability to grow on PCA (Supplementary Fig. S15). These results indicate that the evolved *catB* allele is required to support the PCA-growth phenotype in this background.

**Figure 9.**
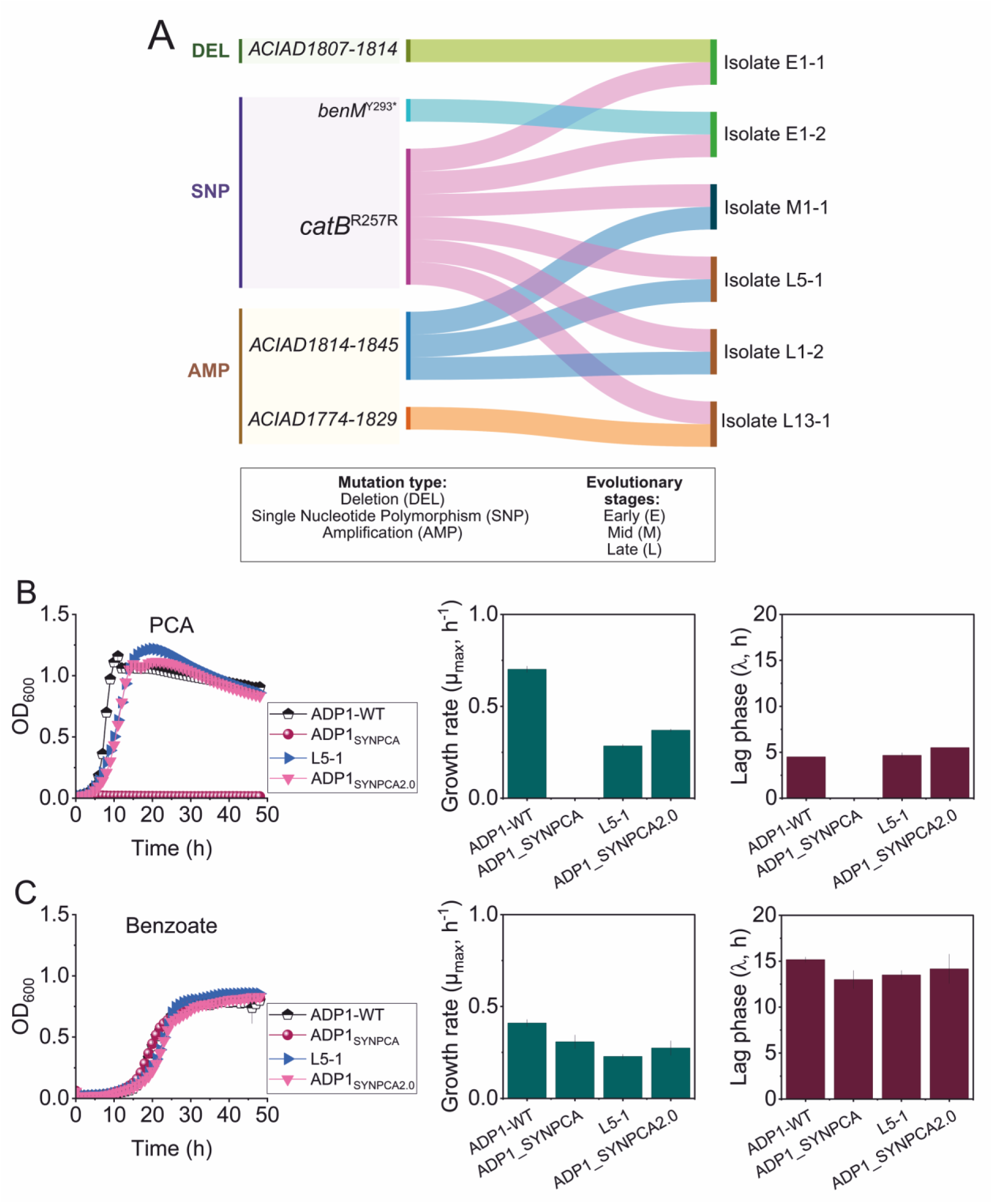
Evaluation and reverse engineering of the ALE-derived mutations. A) Mutations were identified by whole-genome sequencing of the representative isolates (E: early, M: mid, L: late evolutionary stages) (the Sankey diagram was built using SankeyMATIC online tool), detailed description of the mutations are presented in Supplementary Table S8. Growth profile of reverse engineered strains ADP1_SYNPCA2.0_, with the unevolved strain ADP1_SYNPCA_ and the evolved strain L5-1 in MSM with B) 25 mM PCA and C) 10 mM benzoate. μ_max_ represents maximum growth rate and λ represents growth lag. Data represent the mean ± standard deviation of three biological replicates (n = 3). DEL: deletion, SNP: single nucleotide polymorphism, AMP: amplification.

In addition to point mutations, an amplification (∼6-32 copies) was identified in mid and late evolutionary stages (ACIAD1774-1829 and ACIAD1814-1845), with overlapping region (Supplementary Fig. S16). The amplified region encompasses a cluster of poorly characterized genes, including multiple hypothetical proteins, putative membrane-associated proteins, a signal peptide, and a transcriptional regulator (Supplementary Table S9). Because amplification of ACIAD1814-1845 was consistently detected in evolved strains, we further evaluated the functional significance of this region. We deleted the entire ACIAD1814-ACIAD1845 region in five out of six ADP1_SYNPCA_ L strains (L5-1, 1 L6-1, L1-2, L2-2, and L4-2) and assessed whether the removal of this region impacts growth on different substrates. Strains lacking the amplified region were able to grow on PCA as the sole carbon source, but with prolonged exponential phase compared to the controls (Supplementary Fig. S17). In contrast, growth on other carbon sources (benzoate and succinate) showed no pronounced differences. Together, these results indicate that this amplification is not essential for PCA utilization, yet it provides a fitness advantage.

We then performed reverse experiment, introducing the *catB*^R257R;^(CGA→AGA) mutation into the parent strain ADP1_SYNPCA_. First, the *catB* mutated region was PCR-amplified from the evolved isolate. The ADP1_SYNPCA_ strain was then cultured in 500 µL medium with or without transforming DNA for 6 h, followed by centrifugation and transfer of 100 µL into MSM supplemented with 25 mM PCA for overnight incubation. The next day, 10 µL of culture was inoculated into 190 µL of fresh medium in a microplate reader, and growth was monitored over 24 h. As we expected, only cultures receiving the transforming DNA exhibited growth, confirming the functional benefit of the *catB*^R257R;(CGA→AGA)^ mutation (Supplementary Fig. S18). Resulting colonies were streaked onto LB plates containing antibiotics, and selected isolates were grown in MSM supplemented with 25 mM PCA. Whole-genome sequencing confirmed the presence of the targeted catB^R257R;(CGA→AGA)^ mutation. In addition, a reoccurring spontaneous amplification spanning ACIAD1814-1845 was observed.

The reverse-engineered strain, designated as ADP1_SYNPCA2.0_, was then tested for growth in MSM with PCA alongside the ADP1-WT, the evolved strain (L5-1), and the parental strain ADP1_SYNPCA_. Growth comparable to the evolved strain confirmed the functional impact of the *catB*^R257R;(CGA→AGA)^ mutation in PCA utilization. The reverse engineered and evolved strains (L5-1) have lower growth rate on PCA as sole carbon source (μ_max_ = 0.28-0.37 h^-1^) compared to ADP1-WT (μ_max_ = 0.70 h^-1^). Similarly, lower growth rate was also observed on benzoate as sole carbon source (μ_max_ = 0.22-0.31 h^-1^ vs 0.41 h^-1^) (Fig. 6B and 6C). All strains have similar OD_600__max 1.11-1.21 in PCA and 0.81-0.86 in benzoate. Motivated by this result, we next examined the effect of deleting *catB* in the ADP1_SYNPCA_ strain. We speculated that the deletion of *catB* could also induce the PCA utilization through the synthetic protocatechuate branch. However, deletion of *catB* did not mimic the synonymous mutation: Δ*catB* supported growth on 25 mM PCA, but at a lower growth rate than *catB*^R257R;(CGA→AGA)^ (Supplementary Fig. S19).

### Introducing the synthetic pathway with *catB*^R257R;(CGA→AGA)^ mutation in genetically stable ISx_crsA_

Following the successful construction and adaptive evolution in the ADP1_SYNPCA_ strain, we next turned to the genetically stable chassis ISx to validate the functionality of the streamlined PCA utilization pathway within the dual-chassis framework. This design further allowed us to test whether beneficial mutations identified in the complex wild-type background remain effective when transferred into a simplified and genetically stabilized host. ISx_crsA_ strain was chosen over ISx due to its improved growth performance across diverse carbon sources (Supplementary Fig. S3-S5). A synthetic protocatechuate pathway that only contains key essential genes was transformed into ISx_csrA_ ΔPCA ΔDCA into chromosomal location I, previously validated for reliable and tunable expression. The strain was designated as ISx_SYNPCA_. The RAMSES method was subsequently employed to introduce the *catB*^R257R;(CGA→AGA)^ mutation into ISx_SYNPCA_ (Supplementary Fig. S20), previously associated with improved PCA catabolism in evolved strains. All constructs were verified by whole genome sequencing.

The resulting strain (ISx_SYNPCA2.0_) was tested for growth on minimal medium supplemented with PCA as the sole carbon source. While the ISx_SYNPCA_ strain lacking the mutation exhibited no detectable growth, the engineered strain carrying the *catB* ^R257R;(CGA→AGA)^ mutation successfully grew on PCA, reaching final OD_600_ values comparable to those of the ALE-derived isolates (Figure 10). Overall, these findings indicate that the *catB*^R257R;(CGA→AGA)^ is sufficient to restore PCA utilization in a stable genome background when coupled with minimal pathway reconstruction. Furthermore, the *catB^R257R;(CGA→AGA)^* mutation retained its functionality despite the absence of additional genomic elements potentially present in the evolved strain, underscoring its portability and robustness.

**Figure 10.**
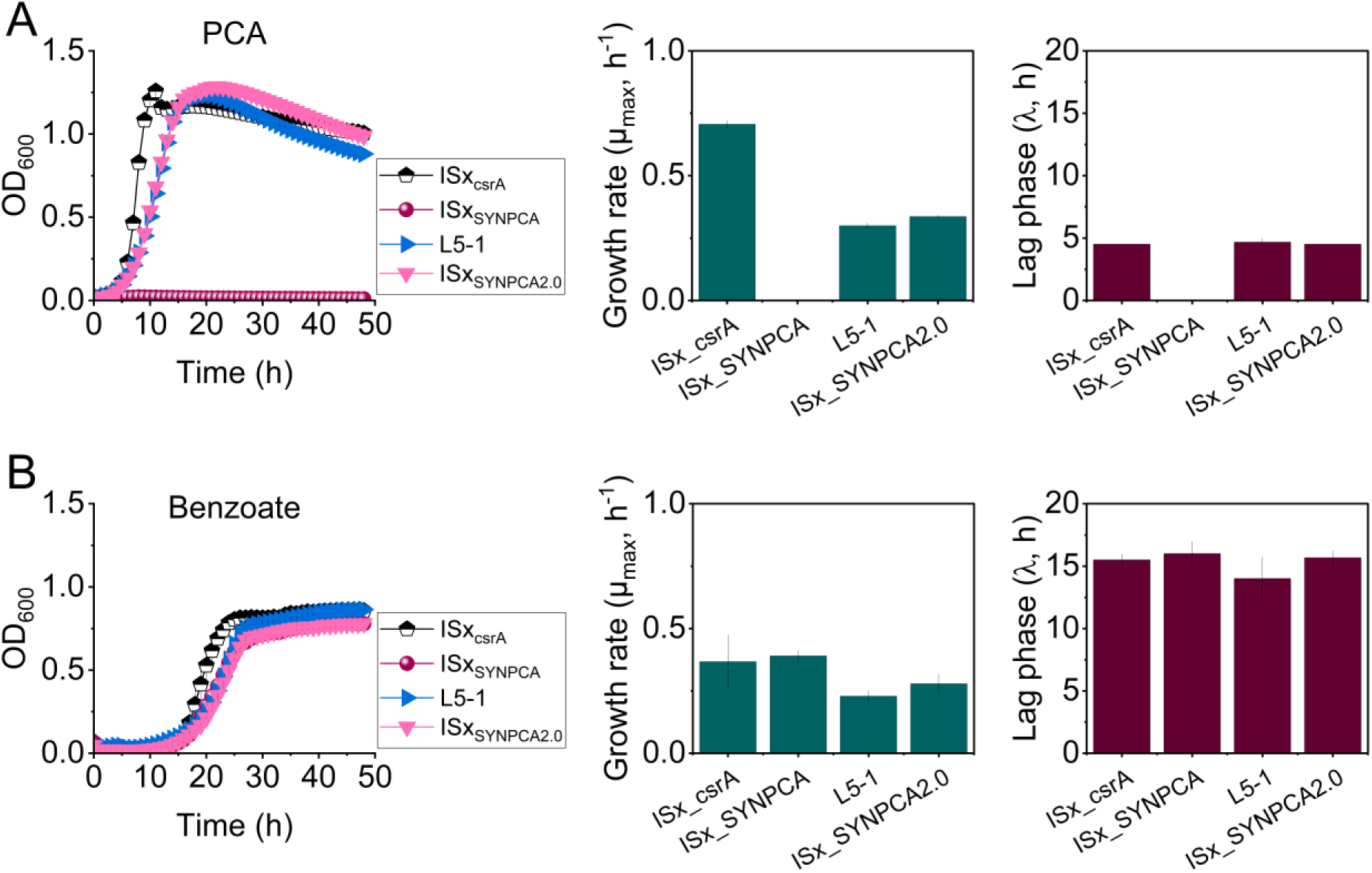
Growth profiles of ISx_SYNPCA_ and ISx_SYNPCA2.0_, with the evolved strain L5-1 in MSM with 25 mM PCA (A) and 10 mM benzoate (B). μ_max_ represents maximum growth rate and λ represents the length of the lag phase in hours. Data represent the mean ± standard deviation of three biological replicates (n = 3).

We further tested the ADP1_SYNPCA2.0_ and ISx_SYNPCA2.0_ strains to utilize mixed substrates, benzoate and PCA. In these experiments with 2 mM benzoate and 2 mM PCA, the control strains ADP1_SYNPCA_ and ISx_SYNPCA_ utilized benzoate but failed to utilize PCA. Notably, strains generated by the dual-chassis strategy (ADP1_SYNPCA2.0_ and ISx_SYNPCA2.0_) were able to consume both benzoate and PCA by the single integrated pathway, although benzoate consumption was expectedly faster (Supplementary Figure S21). Thus, the introduced synthetic pathway accompanied with the identified mutation restored functional catabolism of both substrates, demonstrating that a minimal pathway is sufficient to support not only protocatechuate, but also the utilization of mixed substrates belonging to different branches. In addition, this functional robustness was achieved despite reduction of protocatechuate pathway from ten genes (*pcaUBCDFGHIJK*) in the wild-type strain to a four-gene (*pcaBCGH*) architecture in the refactored strain.

## DISCUSSION

Although pathway streamlining can simplify metabolic networks, rational minimization alone is often insufficient to restore optimal function. Here, we establish a dual-chassis framework that combines rational pathway streamlining with evolutionary adaptation to redesign branched metabolism. Using *Acinetobacter baylyi* ADP1 and its genome-stabilized derivative ISx, we show that evolution-optimized and rationally minimized hosts can provide complementary advantages during pathway reconstruction. As a proof of concept, we dissected the β-ketoadipate pathway (KAP), a central metabolic hub for aromatic compounds degradation (26), to demonstrate how rational simplification coupled with adaptive evolution can be used to restructure and functionally recover a branched metabolic network.

As the initial step, we deleted specific branches, either the catechol (CAT), the protocatechuate (PCA), or the dicarboxylic acids (DCA) branch. This strategy enabled us to isolate the function of each route without interference from parallel catabolic activities. The resulting strains displayed clear specialization: the catechol-branch deletion (ΔCAT) could not utilize coumarate, ferulate, 4HB, or vanillate but readily metabolized benzoate, whereas the protocatechuate-branch deletion (ΔPCA) showed the opposite phenotype. Similarly, deletion of dicarboxylic acid (ΔDCA) branch did not affect the utilization of substrates metabolized via the catechol or protocatechuate branches (Fig. 2, Supplementary Figure S3-S5). These outcomes confirm that these branches operate as semi-independent modules, each capable of sustaining growth on its cognate substrates in isolation.

We also compared the deletion and performance of three different ADP1 strains, showing how stable genome background combined with pathway streamlining affects the growth on aromatic substrates. Similar to a previous report by Gifford et al., 2025 (40), our results also show that ADP1 and ISx_csrA_ outperformed the ISx strain when vanillate served as the sole carbon source. This discrepancy can be explained by the presence of a truncation of a global regulator (*csrA*) in the ISx_csrA_ strain which is involved in the regulation of carbon storage and catabolism (40, 46). CsrA is a global post-transcriptional regulator that directly binds 457 RNAs in *E. coli* and affects the translation of more than 1000 genes, including many regulators of cellular metabolism (46). Previous studies have linked CsrA to the regulation of central carbon metabolism, including glycolysis and glycogenesis (47). The observed phenotype may therefore suggest a role for CsrA in regulating genes involved in vanillate utilization, although such regulatory connections remain to be investigated.

Having established the individual contribution of each branch through targeted deletions, we next asked how pathway partitioning influences substrate utilization at the system level. In ADP1, benzoate is consumed in preference to 4HB, reflecting cross-regulation between the catechol and protocatechuate branches of the β-ketoadipate pathway. Previous studies suggest that this hierarchy is mediated in part by the benzoate-responsive regulators BenM and CatM, which, in the presence of *cis,cis*-muconate (CCM), can repress expression of genes associated with 4HB uptake and protocatechuate branch function (28). Consistent with this regulatory model, physically separating the catechol and protocatechuate branches across strains shifted substrate utilization from sequential to concurrent, as the co-culture consumed benzoate and 4HB simultaneously (Fig. 4B). A similar concept was demonstrated by Singh *et al.* (2019), who engineered two ADP1 strains specialized for benzoate or 4HB consumption through deletion of *pcaGH* or *benD*, respectively (43). However, whereas such single-gene knockouts are designed for a specific substrate pair, our streamlining strategy removes entire redundant catabolic modules, thus avoiding any known or hidden regulatory issues associated with the pathways. Our strategy generated strains that are not only functionally specialized but also genetically more stable and potentially reusable across different pathway configurations regardless of any potentially accumulating pathway intermediates. More broadly, these results suggest that regulatory bottlenecks in branched aromatic catabolism can be overcome not only through targeted pathway blocking, but also through rational genome reduction, which may provide more versatile chassis for the design of synthetic consortia.

These findings also prompted a complementary question: whether the advantages of pathway simplification could be recapitulated within a single engineered strain. To address this, we implemented a dual-chassis system in which the wild-type background (ADP1) was used for pathway construction and adaptive evolution, while the genome-stabilized host (ISx) was reserved for subsequent transfer and validation of evolved mutations. We first constructed a synthetic protocatechuate module resulting in strain ADP1_SYNPCA_ in which the lower-pathways genes of protocatechuate catabolism was rewired to depend on the catechol lower-pathway genes *catDIJF* (Fig. 6A). The pathway, containing the genes *pcaHGBC,* was placed under synthetic regulatory control using an inducible promoter and a strong ribosome binding site, thereby bypassing native transcriptional regulation. This design replaced the native parallel lower-pathway organization with a functionally coupled configuration, allowing us to directly test whether PCA-derived carbon could be sustained through a single shared downstream route.

Our synthetic protocatechuate module, however, did not directly support growth on PCA as sole carbon source in the constructed strain ADP1_SYNPCA_ (Fig. 6C), indicating that the minimal pathway alone could not sustain this function. This suggests that successful pathway installation requires additional genetic, regulatory, or metabolic features beyond the core enzymatic steps. Similar limitations have been previously observed in attempts to rewire *A. baylyi* ADP1 for aromatic catabolism. Baugh et al. (2025) found that introduction of a foreign PCA 2,3-cleavage pathway failed to confer growth on PCA or 4HB unless extensive gene amplification and adaptive mutations occurred (23). More broadly, this is consistent with the challenge of refactoring native pathways, in which removal of native regulation and reorganization into synthetic architectures can introduce secondary constraints not evident from pathway composition alone (48). Together, these results show that streamlined modules are not always functional on their own but depend on proper regulatory and metabolic integration. At the same time, the inability of the minimal strains to grow provided a useful starting point for adaptive laboratory evolution, which restored growth through compensatory changes.

In response to its growth deficiency, the ADP1_SYNPCA_ strain was subjected to adaptive laboratory evolution (ALE) on PCA for two weeks (Fig 7B). During ALE, no mutations were detected within the synthetic protocatechuate pathway itself. Instead, both sequenced lineages acquired the same synonymous mutation in *catB* ^R257R;(CGA→AGA)^, which restored the growth on PCA (Fig 9A, Supplementary Table S8). This repeated evolutionary outcome indicates that the main limitation of the rewired system was not the catalytic capacity of the engineered pathway, but rather to its regulatory network. Although the *catB*^R257R;(CGA→AGA)^ mutation does not alter the encoded amino acid, synonymous substitutions can still influence gene function and fitness through effects on transcription, translation, or expression dynamics (49–51). We therefore hypothesize that the evolved *catB* mutation may indirectly enhance activation of CatM, the LysR-type regulator controlling expression of the cat operon. CatM normally responds to *cis,cis*-muconate to activate cat gene expression (52), and previous work in ADP1 has shown that *catB* mutations can increase CatM-dependent transcription by altering intracellular *cis,cis*-muconate levels (53). However, because our system evolved on PCA rather than benzoate, the relevant activating signal is likely different and remains to be determined. In particular, a potential role of β-carboxymuconate, an intermediate of the protocatechuate branch, warrants further investigation.

Moreover, genome sequencing of evolved strains revealed large segmental amplification (∼29 kb and ∼58 kb), encompassing mainly hypothetical proteins along with several transporters and regulators. Because these amplifications were absent from the early-stage evolved strains (Supplementary Table S7), they likely represent secondary adaptive events acquired during continued evolution rather than the primary basis for restored growth. Although the specific functions of the amplified genes remain unclear, their repeated occurrence suggests that increased dosage of one or more loci may have provided an additional fitness advantage, potentially by improving expression capacity or buffering regulatory constraints imposed by the reduced pathway architecture. Together, these adaptive outcomes emphasize that successful pathway minimization depends not only on preserving core metabolic function, but also on maintaining a host context capable of supporting that simplified architecture.

Transfer of the synthetic pathway together with the evolved mutation (*catB*^R257R;(CGA→AGA)^) into ISx ΔPCA ΔDCA, generating ISx_SYNPCA2.0_, enabled efficient growth on both PCA and benzoate. This result demonstrates that the evolutionary solutions restoring metabolic robustness in ADP1_SYNPCA_ remained functional upon transfer to another host context, supporting their portability beyond the native evolutionary context in which they emerged. Notably, ISx_SYNPCA2.0_ accumulated no additional mutations, consistent with the IS-free nature of the ISx chassis, which minimizes transposition-driven genomic instability and provides a stable environment for pathway deployment (11, 25, 54). Together, these findings highlight the core advantage of the DUET framework: adaptive evolution can be exploited in a flexible chassis to uncover robust functional solutions, which can then be deployed unchanged into a genetically stable and more controllable host for reliable pathway implementation.

These results not only validate the dual-chassis strategy but also define minimal aromatic catabolic architectures that are both biologically functional and genetically tractable. Efficient channelling of diverse lignin-derived aromatics is a central requirement for microbial lignin upgrading, yet this network is often constrained by redundancy, regulatory entanglement, and uneven pathway utilization (55). Using the β-ketoadipate pathway as a case study, we show that simplifying complex native networks can improve substrate routing while revealing the minimal functions required for the assimilation of lignin-related aromatics.

In conclusion, this work shows that the branched aromatic degradation network of *A. baylyi* ADP1 can be systematically reduced and functionally rebuilt using a combination of rational design and adaptive evolution. We first simplified the native network by deleting pathway branches, then reconstructed a minimal synthetic route that merges protocatechuate and catechol metabolism. Adaptive laboratory evolution in an evolution-competent host restored efficient pathway function, and the identified beneficial mutations were subsequently transferred to a genomically stable, equally reduced ISx strain. By this systematic approach we demonstrated that aromatic catabolism can be rewired while preserving the capacity to assimilate substrates from both branches with reduced pathway interference.

More broadly, these streamlined genetic backgrounds provide a modular framework for assembling *de novo* synthetic pathways, enabling iterative optimization of enzyme variants or pathway segments from different organisms in an evolution-competent host. Within this framework, ADP1 serves as a versatile platform for pathway discovery, testing, and refinement, after which validated designs can be transferred into more stable or application-oriented hosts. This study therefore establishes – not only a novel platform for lignin valorization – but also a generalizable framework for metabolic reprogramming and for systematic exploration of metabolic network organization.

## ACKNOWLEDGEMENTS

We thank Jin Luo for performing nanopore sequencing and for helpful discussions related to adaptive laboratory evolution, Elena Efimova for providing the HPLC method. We acknowledge the Texas Advanced Computing Center (TACC) at The University of Texas at Austin for providing high performance computing resources. Schematic figures were created with BioRender.com.

## AUTHOR CONTRIBUTIONS

Kesi Kurnia: Conceptualization, Formal analysis, Visualization, Methodology, Validation, Writing—original draft, Writing—review & editing. Isaac Gifford: Methodology, Writing—review & editing. Ville Santala: Conceptualization, Supervision, Funding acquisition, Writing—review & editing. Jeffrey E Barrick: Supervision, Writing—review & editing. Suvi Santala: Conceptualization, Supervision, Funding acquisition, Writing—review & editing.

## SUPPLEMENTARY DATA

Supplementary data is available at NAR online.

## DATA AVAILABILITY

The data that support the findings of this study are included within the article or the additional files. The corresponding author is willing to provide the raw data related to this manuscript upon reasonable request. FASTQ files from genome sequencing of ADP1_SYNPCA_ and ISx_SYNPCA2.0_ are available from the NCBI Sequence Read Archive (PRJNA1470083).

## REFERENCES

1. Xu, X., Meier, F., Blount, B.A., Pretorius, I.S., Ellis, T., Paulsen, I.T. and Williams, T.C. (2023) Trimming the genomic fat: minimising and re-functionalising genomes using synthetic biology. Nat Commun, 14, 1984.

2. Kim, K., Choe, D., Cho, S., Palsson, B. and Cho, B.-K. (2024) Reduction-to-synthesis: the dominant approach to genome-scale synthetic biology. Trends in Biotechnology, 42, 1048–1063.

3. Park, M.K., Lee, S.H., Yang, K.S., Jung, S.-C., Lee, J.H. and Kim, S.C. (2014) Enhancing recombinant protein production with an Escherichia coli host strain lacking insertion sequences. Appl Microbiol Biotechnol, 98, 6701–6713.

4. Martínez-García, E., Nikel, P.I., Aparicio, T. and De Lorenzo, V. (2014) Pseudomonas 2.0: genetic upgrading of P. putida KT2440 as an enhanced host for heterologous gene expression. Microb Cell Fact, 13, 159.

5. Wang, Y.-H., Wei, K.Y. and Smolke, C.D. (2013) Synthetic Biology: Advancing the Design of Diverse Genetic Systems. Annu. Rev. Chem. Biomol. Eng., 4, 69–102.

6. Temme, K., Zhao, D. and Voigt, C.A. (2012) Refactoring the nitrogen fixation gene cluster from *Klebsiella oxytoca*. Proc. Natl. Acad. Sci. U.S.A., 109, 7085–7090.

7. Sandberg, T.E., Salazar, M.J., Weng, L.L., Palsson, B.O. and Feist, A.M. (2019) The emergence of adaptive laboratory evolution as an efficient tool for biological discovery and industrial biotechnology. Metabolic Engineering, 56, 1–16.

8. Gifford, I., Suárez, G.A. and Barrick, J.E. (2024) Evolution recovers the fitness of Acinetobacter baylyi strains with large deletions through mutations in deletion-specific targets and global post-transcriptional regulators. PLoS Genet, 20, e1011306.

9. Choe, D., Lee, J.H., Yoo, M., Hwang, S., Sung, B.H., Cho, S., Palsson, B., Kim, S.C. and Cho, B.-K. (2019) Adaptive laboratory evolution of a genome-reduced Escherichia coli. Nat Commun, 10, 935.

10. Bleem, A.C., Kuatsjah, E., Johnsen, J., Mohamed, E.T., Alexander, W.G., Kellermyer, Z.A., Carroll, A.L., Rossi, R., Schlander, I.B., Peabody V, G.L., et al. (2024) Evolution and engineering of pathways for aromatic O-demethylation in Pseudomonas putida KT2440. Metabolic Engineering, 84, 145–157.

11. Suárez, G.A., Renda, B.A., Dasgupta, A. and Barrick, J.E. (2017) Reduced Mutation Rate and Increased Transformability of Transposon-Free Acinetobacter baylyi ADP1-ISx. Appl Environ Microbiol, 83, e01025–17.

12. Metzgar, D. (2004) Acinetobacter sp. ADP1: an ideal model organism for genetic analysis and genome engineering. Nucleic Acids Research, 32, 5780–5790.

13. Elliott, K.T. and Neidle, E.L. (2011) Acinetobacter baylyi ADP1: Transforming the choice of model organism. IUBMB Life, 63, 1075–1080.

14. Santala, S. and Santala, V. (2021) *Acinetobacter baylyi* ADP1—naturally competent for synthetic biology. Essays in Biochemistry, 65, 309–318.

15. Salmela, M., Lehtinen, T., Efimova, E., Santala, S. and Santala, V. (2019) Alkane and wax ester production from lignin-related aromatic compounds. Biotech & Bioengineering, 116, 1934–1945.

16. Luo, J., Lehtinen, T., Efimova, E., Santala, V. and Santala, S. (2019) Synthetic metabolic pathway for the production of 1-alkenes from lignin-derived molecules. Microb Cell Fact, 18, 48.

17. Kurnia, K., Efimova, E., Santala, V. and Santala, S. (2024) Metabolic engineering of Acinetobacter baylyi ADP1 for naringenin production. Metabolic Engineering Communications, 19, e00249.

18. Arvay, E., Biggs, B.W., Guerrero, L., Jiang, V. and Tyo, K. (2021) Engineering Acinetobacter baylyi ADP1 for mevalonate production from lignin-derived aromatic compounds. Metabolic Engineering Communications, 13, e00173.

19. Liu, C., Juvonen, V., Meriläinen, E., Efimova, E., Luo, J., Salmela, M., Santala, S. and Santala, V. (2025) Cis, cis-muconic acid production from lignin related molecules byAcinetobacter baylyi ADP1. Microb Cell Fact, 24, 150.

20. Jiang, X., Palazzotto, E., Wybraniec, E., Munro, L.J., Zhang, H., Kell, D.B., Weber, T. and Lee, S.Y. (2020) Automating Cloning by Natural Transformation. ACS Synth. Biol., 9, 3228–3235.

21. Maiti, S., Singh, P., Prasad J, V., Muthukrishnan, A.B., Blank, L.M. and Jayaraman, G. (2025) A consortium-based approach to adaptive laboratory evolution of *Acinetobacter baylyi* ADP1 reveals novel genetic targets for lignin valorization. Journal of Applied Microbiology, 136, lxaf138.

22. Luo, J., McIntyre, E.A., Bedore, S.R., Santala, V., Neidle, E.L. and Santala, S. (2022) Characterization of Highly Ferulate-Tolerant Acinetobacter baylyi ADP1 Isolates by a Rapid Reverse Engineering Method. Appl Environ Microbiol, 88, e01780–21.

23. Baugh, A.C., Tumen-Velasquez, M.P., Zempel, I.R., Duscent-Maitland, C.V., Slarks, L.E., Defalco, J.B., Johnson, C.W., Beckham, G.T. and Neidle, E.L. (2025) Rewiring Aromatic Compound Consumption: Chromosomal Amplification and Evolution of a Foreign Pathway in *Acinetobacter baylyi* ADP1. ACS Synth. Biol., 10.1021/acssynbio.5c00341.

24. Tumen-Velasquez, M., Johnson, C.W., Ahmed, A., Dominick, G., Fulk, E.M., Khanna, P., Lee, S.A., Schmidt, A.L., Linger, J.G., Eiteman, M.A., et al. (2018) Accelerating pathway evolution by increasing the gene dosage of chromosomal segments. Proc. Natl. Acad. Sci. U.S.A., 115, 7105–7110.

25. Suárez, G.A., Dugan, K.R., Renda, B.A., Leonard, S.P., Gangavarapu, L.S. and Barrick, J.E. (2020) Rapid and assured genetic engineering methods applied to Acinetobacter baylyi ADP1 genome streamlining. Nucleic Acids Research, 48, 4585–4600.

26. Harwood, C.S. and Parales, R.E. (1996) THE β-KETOADIPATE PATHWAY AND THE BIOLOGY OF SELF-IDENTITY. Annu. Rev. Microbiol., 50, 553–590.

27. Gaines, G.L., Smith, L. and Neidle, E.L. (1996) Novel nuclear magnetic resonance spectroscopy methods demonstrate preferential carbon source utilization by Acinetobacter calcoaceticus. J Bacteriol, 178, 6833–6841.

28. Brzostowicz, P.C., Reams, A.B., Clark, T.J. and Neidle, E.L. (2003) Transcriptional Cross-Regulation of the Catechol and Protocatechuate Branches of the ␤-Ketoadipate Pathway Contributes to Carbon Source-Dependent Expression of the Acinetobacter sp. Strain ADP1 pobA Gene. APPL. ENVIRON. MICROBIOL., 69.

29. Bleichrodt, F.S., Fischer, R. and Gerischer, U.C. (2010) The β-ketoadipate pathway of Acinetobacter baylyi undergoes carbon catabolite repression, cross-regulation and vertical regulation, and is affected by Crc. Microbiology, 156, 1313–1322.

30. Gifford, I., Vergis, M.R. and Barrick, J.E. (2025) Genome evolution of *Acinetobacter baylyi* ADP1 during laboratory domestication: acquired mutations impact competence and metabolism. Appl Environ Microbiol, 91, e00936–25.

31. Tuomela, H., Koivisto, J., Efimova, E. and Santala, S. (2025) Conversion and upgrading of syringate by Acinetobacter baylyi ADP1. Microb Cell Fact, 24, 209.

32. Schuster, L.A. and Reisch, C.R. (2021) A plasmid toolbox for controlled gene expression across the Proteobacteria. Nucleic Acids Research, 49, 7189–7202.

33. Biggs, B.W., Bedore, S.R., Arvay, E., Huang, S., Subramanian, H., McIntyre, E.A., Duscent-Maitland, C.V., Neidle, E.L. and Tyo, K.E.J. (2020) Development of a genetic toolset for the highly engineerable and metabolically versatile Acinetobacter baylyi ADP1. Nucleic Acids Research, 48, 5169–5182.

34. Luo, J., Efimova, E., Volke, D.C., Santala, V. and Santala, S. (2022) Engineering cell morphology by CRISPR interference in *Acinetobacter baylyi* ADP1. Microbial Biotechnology, 15, 2800–2818.

35. Dvořák, P., Burýšková, B., Popelářová, B., Ebert, B.E., Botka, T., Bujdoš, D., Sánchez-Pascuala, A., Schöttler, H., Hayen, H., De Lorenzo, V., et al. (2024) Synthetically-primed adaptation of Pseudomonas putida to a non-native substrate D-xylose. Nat Commun, 15, 2666.

36. Deatherage, D.E. and Barrick, J.E. (2014) Identification of Mutations in Laboratory-Evolved Microbes from Next-Generation Sequencing Data Using breseq. In Sun, L., Shou, W. (eds), Engineering and Analyzing Multicellular Systems, Methods in Molecular Biology. Springer New York, New York, NY, Vol. 1151, pp. 165–188.

37. Wells, T. and Ragauskas, A.J. (2012) Biotechnological opportunities with the β-ketoadipate pathway. Trends in Biotechnology, 30, 627–637.

38. Siehler, S.Y., Dal, S., Fischer, R., Patz, P. and Gerischer, U. (2007) Multiple-Level Regulation of Genes for Protocatechuate Degradation in *Acinetobacter baylyi* Includes Cross-Regulation. Appl Environ Microbiol, 73, 232–242.

39. Fischer, R., Bleichrodt, F.S. and Gerischer, U.C. (2008) Aromatic degradative pathways in Acinetobacter baylyi underlie carbon catabolite repression. Microbiology, 154, 3095–3103.

40. Gifford, I., Vergis, M.R. and Barrick, J.E. (2025) Genome evolution of *Acinetobacter baylyi* ADP1 during laboratory domestication: acquired mutations impact competence and metabolism. Appl Environ Microbiol, 10.1128/aem.00936-25.

41. Parke, D., Garcia, M.A. and Ornston, L.N. (2001) Cloning and Genetic Characterization of *dca* Genes Required for β-Oxidation of Straight-Chain Dicarboxylic Acids in *Acinetobacter* sp. Strain ADP1. Appl Environ Microbiol, 67, 4817–4827.

42. Pardo, I., Jha, R.K., Bermel, R.E., Bratti, F., Gaddis, M., McIntyre, E., Michener, W., Neidle, E.L., Dale, T., Beckham, G.T., et al. (2020) Gene amplification, laboratory evolution, and biosensor screening reveal MucK as a terephthalic acid transporter in Acinetobacter baylyi ADP1. Metabolic Engineering, 62, 260–274.

43. Singh, A., Bedore, S.R., Sharma, N.K., Lee, S.A., Eiteman, M.A. and Neidle, E.L. (2019) Removal of aromatic inhibitors produced from lignocellulosic hydrolysates by Acinetobacter baylyi ADP1 with formation of ethanol by Kluyveromyces marxianus. Biotechnol Biofuels, 12, 91.

44. Santala, S., Efimova, E. and Santala, V. (2018) Dynamic decoupling of biomass and wax ester biosynthesis in Acinetobacter baylyi by an autonomously regulated switch. Metabolic Engineering Communications, 7, e00078.

45. Tuomela, H., Koivisto, J., Efimova, E. and Santala, S. (2025) Conversion and upgrading of S-lignin related syringate by Acinetobacter baylyi ADP1. 10.21203/rs.3.rs-6218493/v1.

46. Potts, A.H., Vakulskas, C.A., Pannuri, A., Yakhnin, H., Babitzke, P. and Romeo, T. (2017) Global role of the bacterial post-transcriptional regulator CsrA revealed by integrated transcriptomics. Nat Commun, 8, 1596.

47. Timmermans, J. and Van Melderen, L. (2009) Conditional Essentiality of the *csrA* Gene in *Escherichia coli*. J Bacteriol, 191, 1722–1724.

48. Smanski, M.J., Bhatia, S., Zhao, D., Park, Y., B A Woodruff, L., Giannoukos, G., Ciulla, D., Busby, M., Calderon, J., Nicol, R., et al. (2014) Functional optimization of gene clusters by combinatorial design and assembly. Nat Biotechnol, 32, 1241–1249.

49. Bailey, S.F., Hinz, A. and Kassen, R. (2014) Adaptive synonymous mutations in an experimentally evolved Pseudomonas fluorescens population. Nat Commun, 5, 4076.

50. Agashe, D., Sane, M., Phalnikar, K., Diwan, G.D., Habibullah, A., Martinez-Gomez, N.C., Sahasrabuddhe, V., Polachek, W., Wang, J., Chubiz, L.M., et al. (2016) Large-Effect Beneficial Synonymous Mutations Mediate Rapid and Parallel Adaptation in a Bacterium. Mol Biol Evol, 33, 1542–1553.

51. Kristofich, J., Morgenthaler, A.B., Kinney, W.R., Ebmeier, C.C., Snyder, D.J., Old, W.M., Cooper, V.S. and Copley, S.D. (2018) Synonymous mutations make dramatic contributions to fitness when growth is limited by a weak-link enzyme. PLoS Genet, 14, e1007615.

52. Craven, S.H., Ezezika, O.C., Haddad, S., Hall, R.A., Momany, C. and Neidle, E.L. (2009) Inducer responses of BenM, a LysR-type transcriptional regulator from *Acinetobacter baylyi* ADP1. Molecular Microbiology, 72, 881–894.

53. Cosper, N.J., Collier, L.S., Clark, T.J., Scott, R.A. and Neidle, E.L. (2000) Mutations in *catB*, the Gene Encoding Muconate Cycloisomerase, Activate Transcription of the Distal *ben* Genes and Contribute to a Complex Regulatory Circuit in *Acinetobacter* sp. Strain ADP1. J Bacteriol, 182, 7044–7052.

54. Consuegra, J., Gaffé, J., Lenski, R.E., Hindré, T., Barrick, J.E., Tenaillon, O. and Schneider, D. (2021) Insertion-sequence-mediated mutations both promote and constrain evolvability during a long-term experiment with bacteria. Nat Commun, 12, 980.

55. Liu, R.-Y., Li, B.-Z., Yuan, Y.-J. and Liu, Z.-H. (2025) Multiscale metabolic engineering in biological lignin valorization. The Innovation, 6, 100993.

