## Supplementary Figures for "Dual-Chassis Strategy for Bridging Adaptive Evolution and Rational Design for Synthetic Biology"

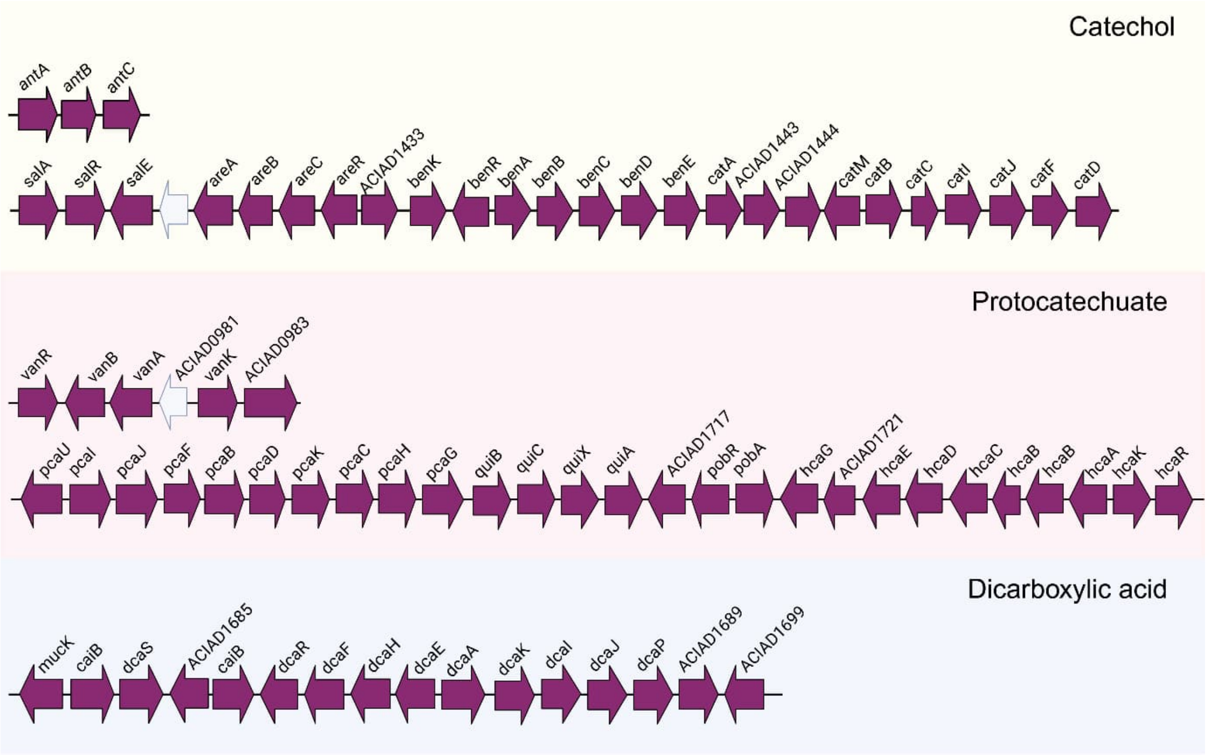


Supplementary Figure S1. Genes are responsible for aromatic degradation pathway in ADP1. The details of deleted regions and their annotation can be found in Supplementary Table S5.

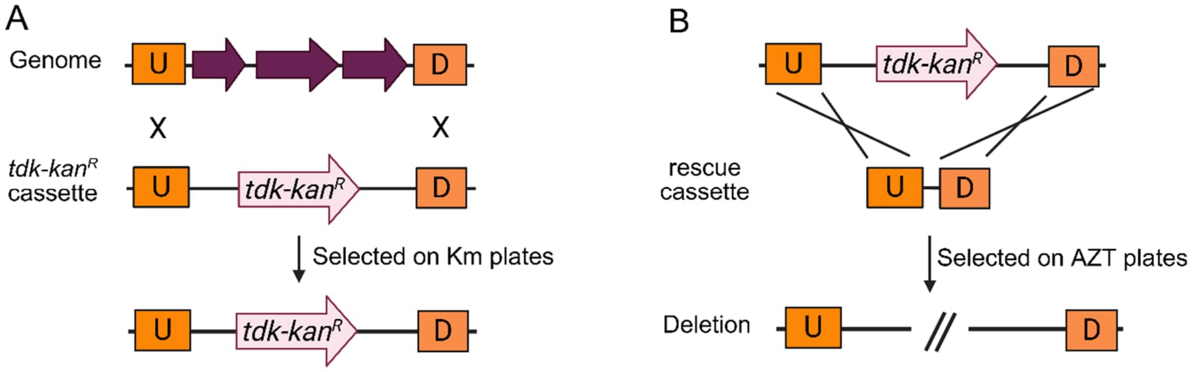


Supplementary Figure S2. Schematic representation of constructing deletion strain of ADP1 using Golden Gate transformation (1). A) The target locus was replaced with a *tdk-kanR* cassette flanked by ~1 kb upstream and downstream homology regions to enable homologous recombination, and successful integrants were selected on kanamycin plates, B) The rescue cassette was then integrated at the same locus to excise the *tdk-kanR* marker, with colonies selected on AZT-containing agar plates. U; upstream, D; downstream.

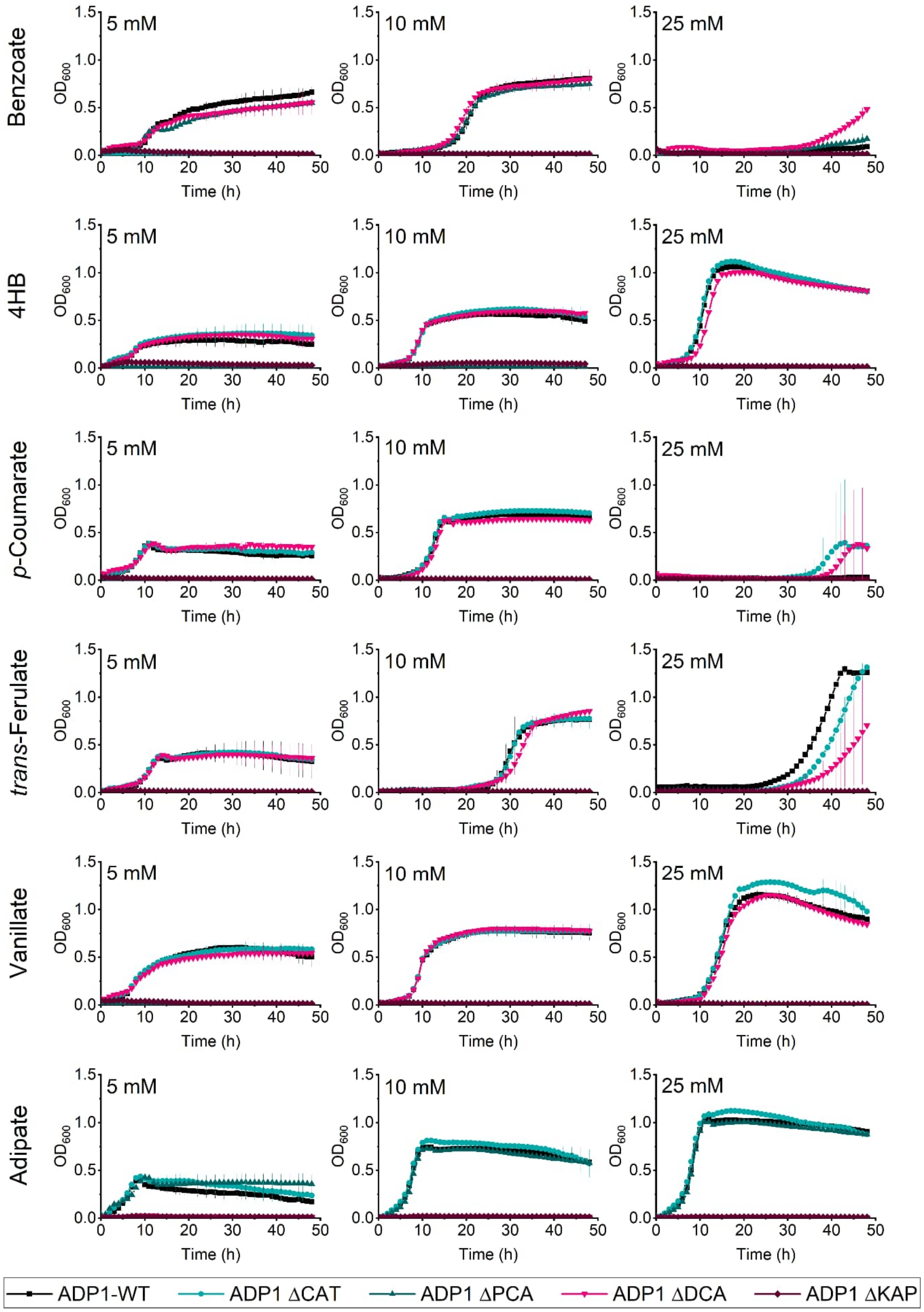


Supplementary Figure S3. Growth profile on ADP1 and derived strains on different substrates. Data points represent the means ± standard deviations calculated from three (n=3) biological replicates. Raw data are available in Supplementary Data 1.

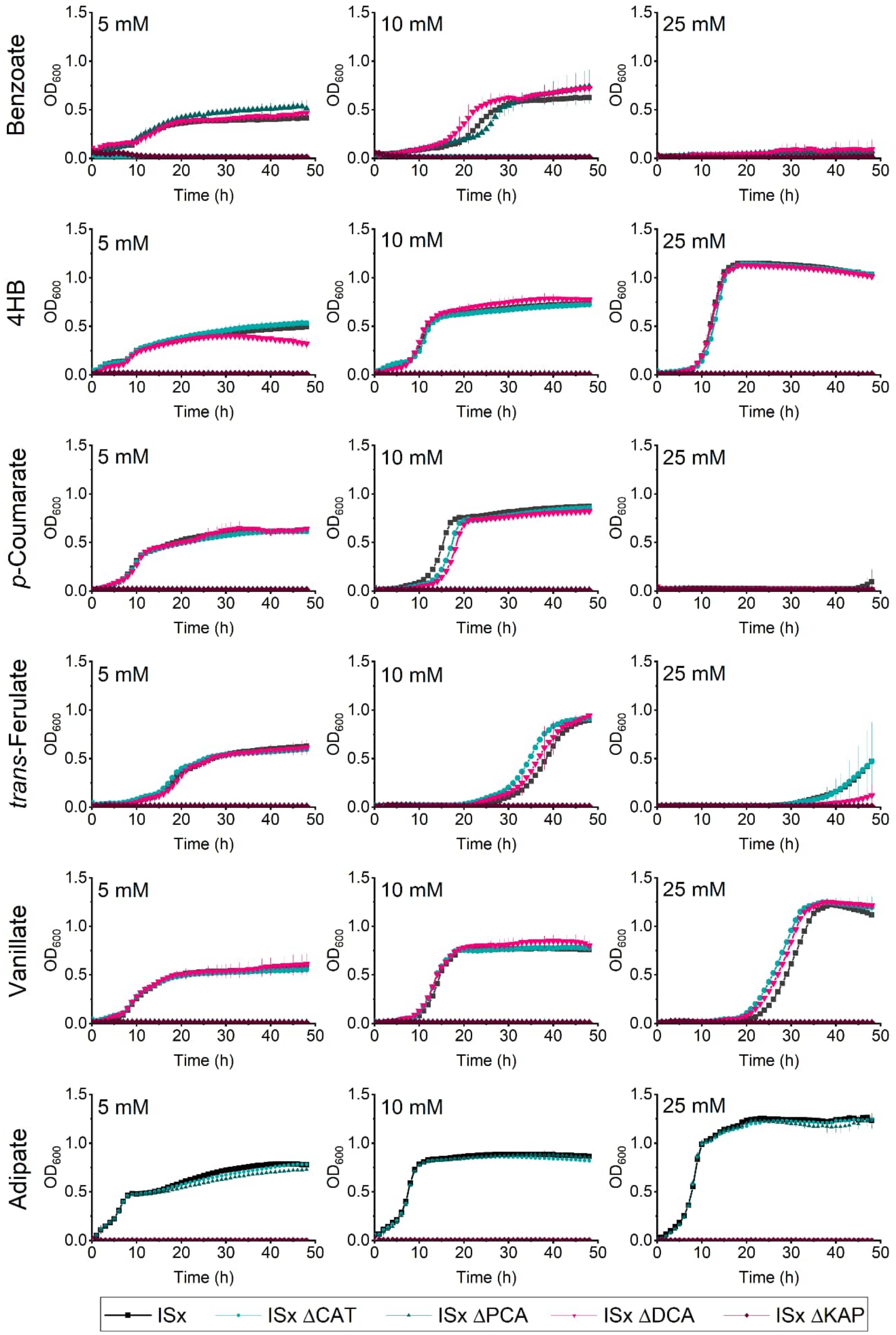


Supplementary Figure S4. Growth profile on ISx and derived strains on different substrates. Data points represent the means ± standard deviations calculated from three (n=3) biological replicates. Raw data are available in Supplementary Data 1.

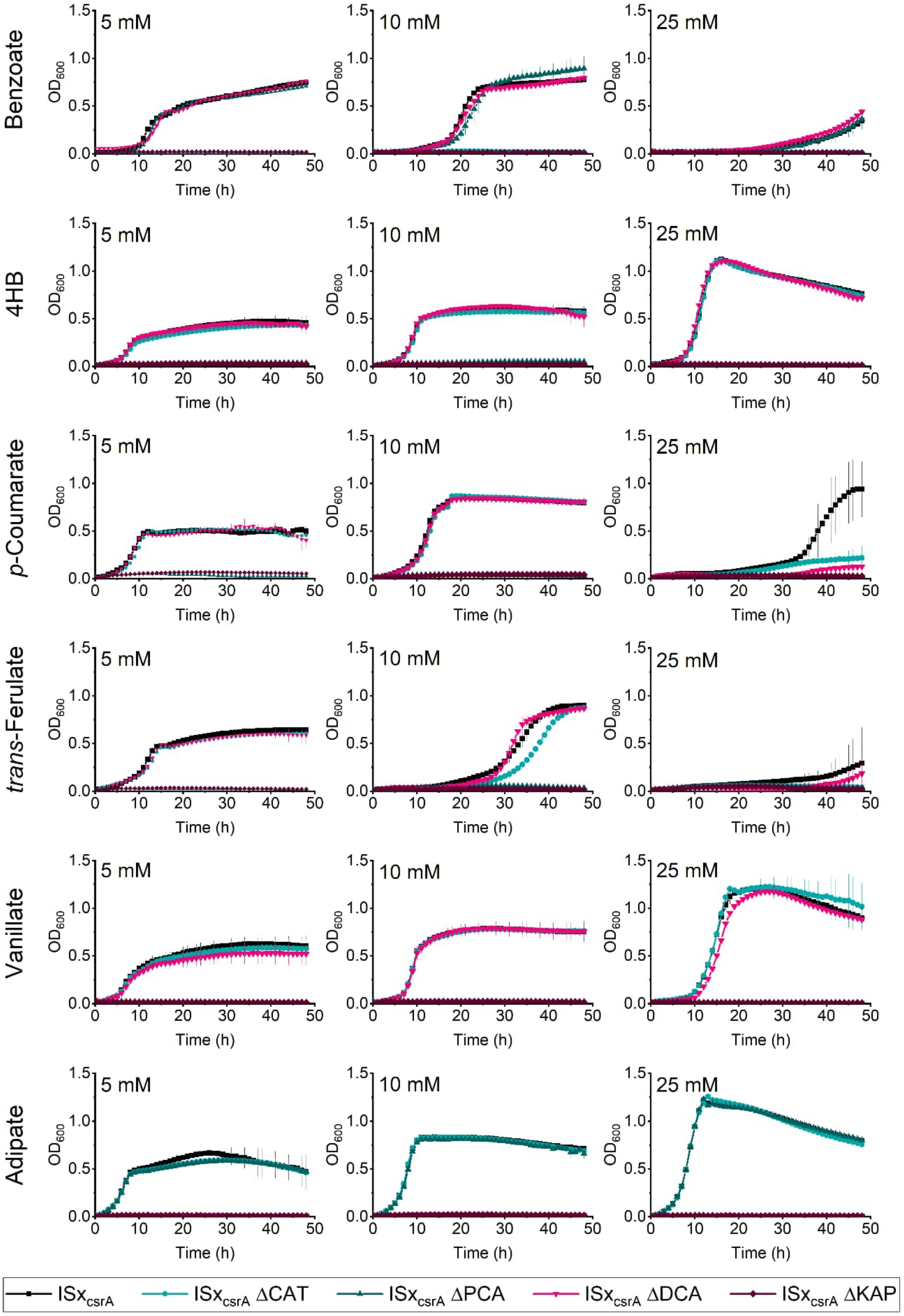


Supplementary Figure S5. Growth profile on $ISx_{csrA}$ and derived-strains on different substrates. Data points represent the means ± standard deviations calculated from three (n=3) biological replicates. Raw data are available in Supplementary Data 1.

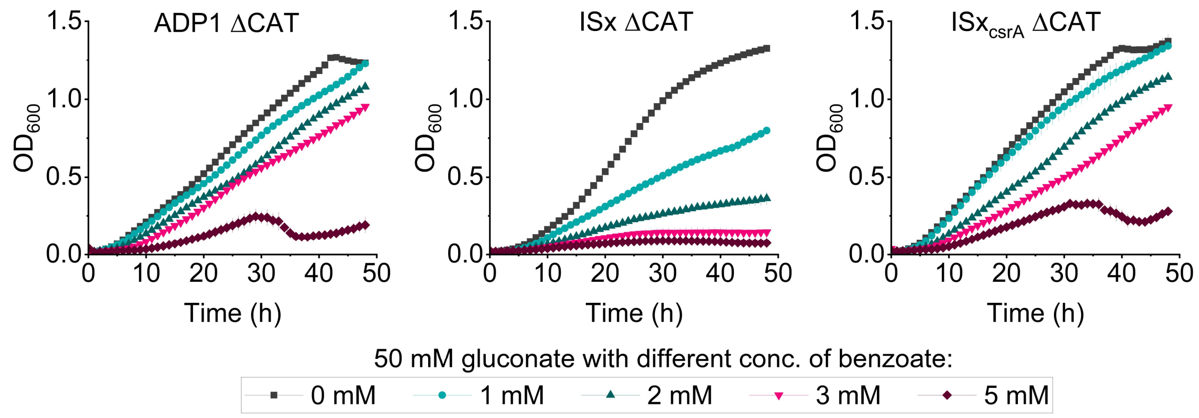


Supplementary Figure S6. Growth of catechol deletion strains (ΔCAT) in MSM with 50 mM gluconate and different concentration of benzoate. Data points represent the means ± standard deviations calculated from three (n=3) biological replicates.

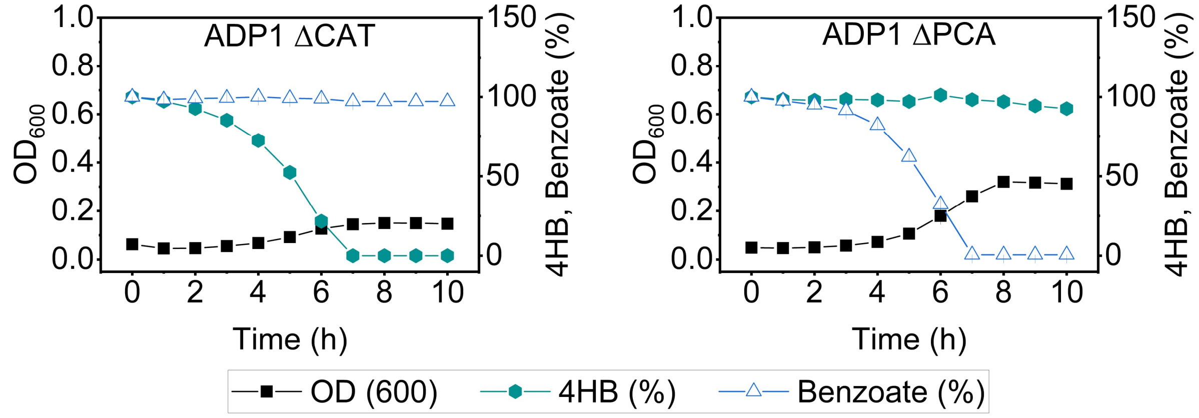


Supplementary Figure S7. $OD_{600}$ values and substrates consumption of monoculture ADP1 ΔCAT (A) and ADP1 ΔPCA (B). Cells were grown in MSM with 2 mM of 4HB and 2 mM of benzoate. Data points represent the means ± standard deviations calculated from three (n=3) biological replicates.

### Growth and Fluorescence measurement

Two chromosomal integration sites were evaluated: the *poxB* locus (site I; ADP1 mRFP-L1) and the *pca-qui-pob-hca* cluster (site II). At site II, two mRFP strains were constructed with the integration cassette inserted in opposite orientations: ADP1 mRFP-L2 in the forward orientation and ADP1 mRFP-L2R in the reverse orientation. While one orientation supported normal growth, the other exhibited poor growth in both LB and MSM supplemented with 50 mM gluconate and 0.2% (w/v) casamino acids. In 5 mL liquid cultures incubated for 24 h in triplicate, no measurable growth was observed, and the strain also displayed slow colony formation on agar plates. Due to this impaired growth during pre-culture, the strain was excluded from further analysis.

Growth of ADP1 mRFP strains was assessed in 5 mL of medium containing 50 mM gluconate, 0.2% (w/v) casamino acids, and Km. Cultures were inoculated at an initial $OD_{660}$ of 0.1 and incubated at 30 °C with shaking. Inducers were added when cultures reached an $OD_{660}$ of 0.2-0.3, at final concentrations ranging from 0 to 100 µM. Prior to measurement, cells were washed twice with 1× PBS and resuspended in the same buffer. $OD_{660}$ and mRFP fluorescence were measured at 20 h post-induction using a Spark microplate reader (Tecan, Switzerland) with the gain set to 40. Measurements were performed in triplicate. Fluorescence values were background-subtracted using blank controls and normalized to cell density (mRFP/$OD_{660}$). Data are presented as mean ± standard deviation (s.d.).

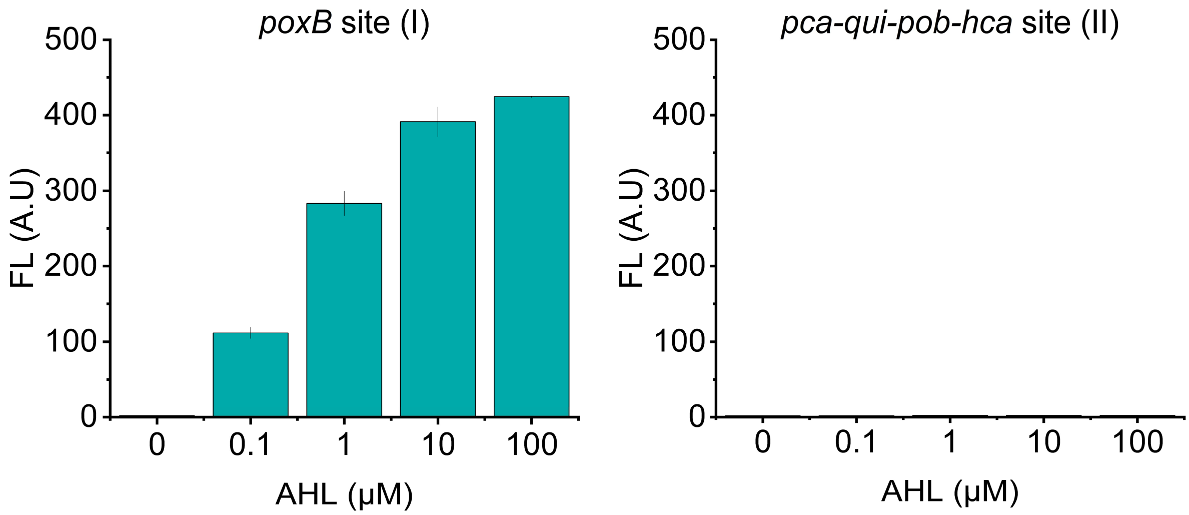


Supplementary Figure S8. Fluorescence assay of ADP1 with genomic integration of mRFP at two different locations: 1) ADP1 mRFP-L1 in the *poxB* site (I) and 2) ADP1 mRFP-L2 in the *pca-qui-pob-hca* site (II). The ADP1 mRFP-L2R strain with reverse genomic integration of mRFP to *pca-qui-pob-hca* site (II) failed to grow in both LB and MSM supplemented with 50 mM gluconate and 0.2% (w/v) casamino acids when cultured in 5 mL volumes for 24 h in triplicate. As the strain did not grow during pre-culture, no further tests were performed, and it was excluded from this analysis. Data points represent the means ± standard deviations calculated from three biological replicates (n=3).

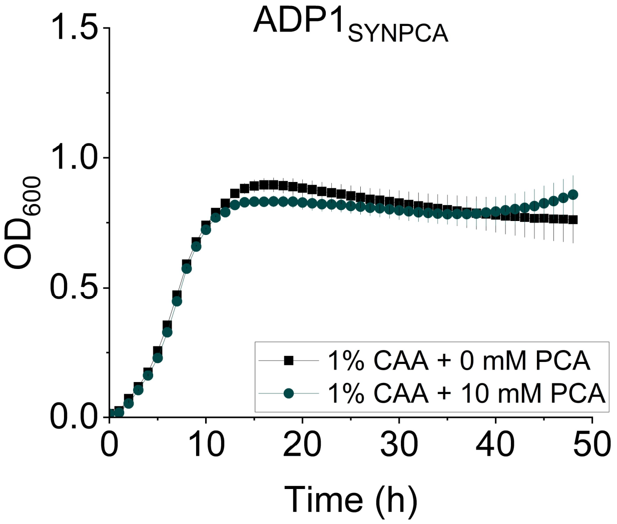


Supplementary Figure S9. Growth profile of ADP1$_{SYNPCA}$ strain in minimal salts medium (MSM) supplemented with 1% (w/v) casamino acids, with or without 10 mM PCA. Data points represent mean values ± standard deviations from three biological replicates (n = 3).

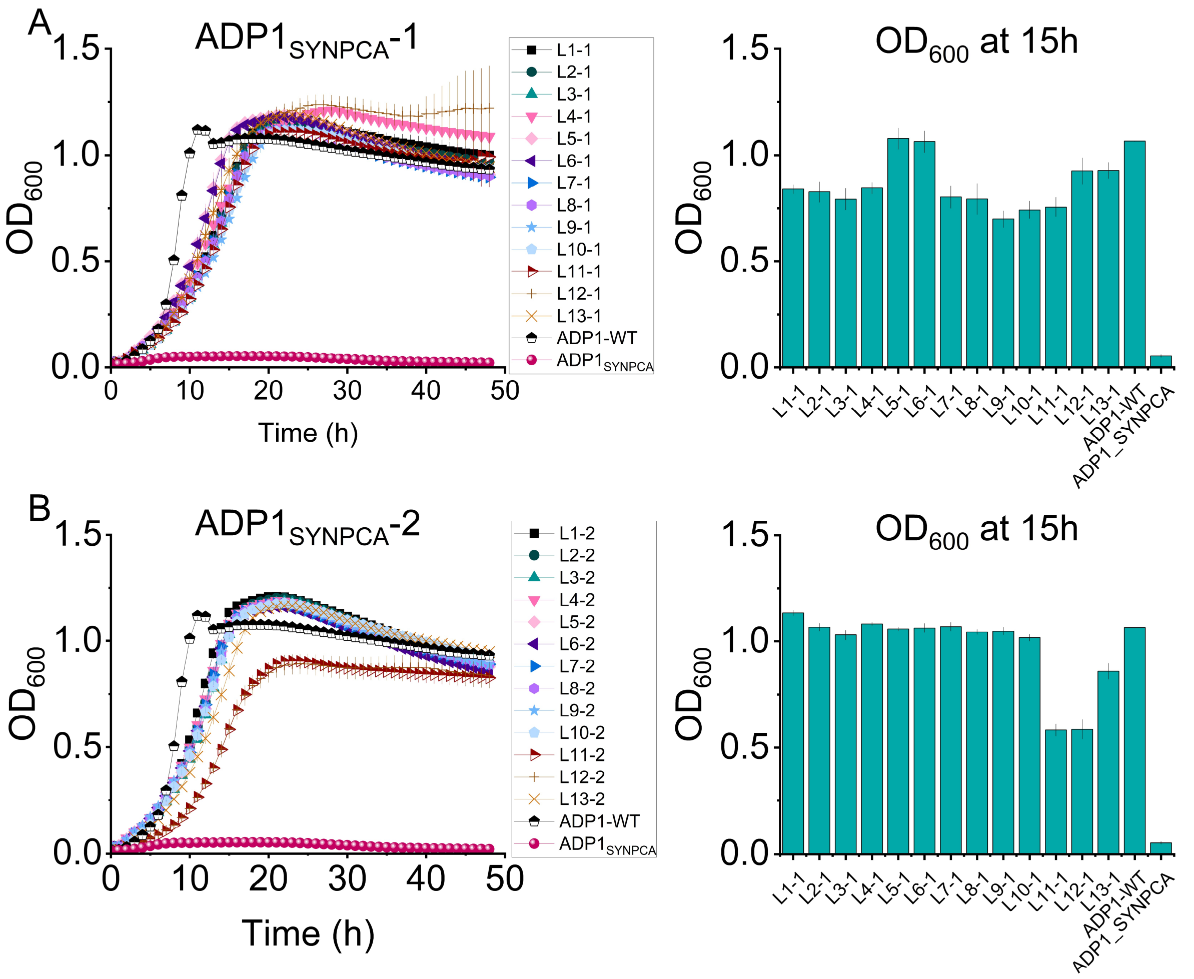


Supplementary Figure S10. Growth profile and $OD_{600}$ values at 15h of L strains from two evolutionary lines (A) $ADP1_{SYNPCA}$-1 and (B) $ADP1_{SYNPCA}$-2. Cells were grown in MSM with 25 mM PCA. Data points represent the means ± standard deviations calculated from three biological replicates (n=3).

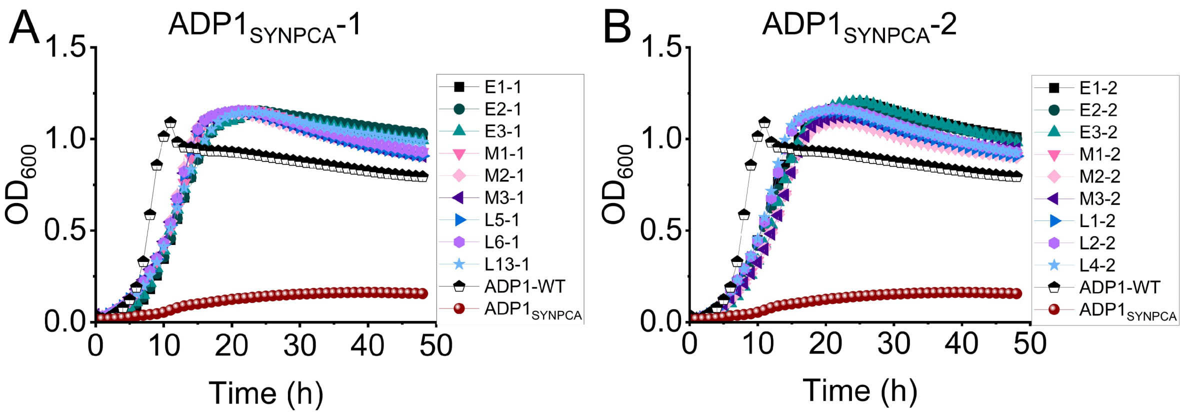


Supplementary Figure S11. Growth profiles of evolved strains (E, M, and L) from two evolutionary lines (A) ADP1$_{SYNPCA}$-1 and (B) ADP1$_{SYNPCA}$-2 in MSM with 25 mM PCA with Km and 10 μM of AHL. Data points represent the mean ± standard deviation from *n* = 3 biological replicates. Underlying data and calculated growth parameters are provided in Supplementary Data 2.

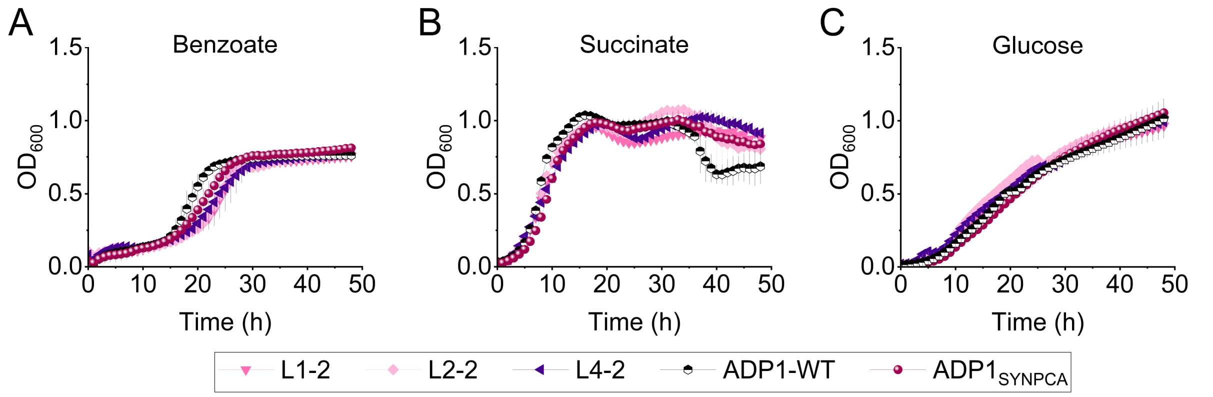


Supplementary Figure S12. Growth profiles of the late evolved strains (L) from evolutionary line 2 compared to ADP1-WT and parental strain ADP1$_{SYNPCA}$ in MSM supplemented with different carbon sources: A) 10 mM benzoate, B) 50 mM succinate, C) 50 mM glucose. Data points represent the mean ± standard deviation from *n* = 3 biological replicates. Underlying data and calculated growth parameters are provided in Supplementary Data 3.

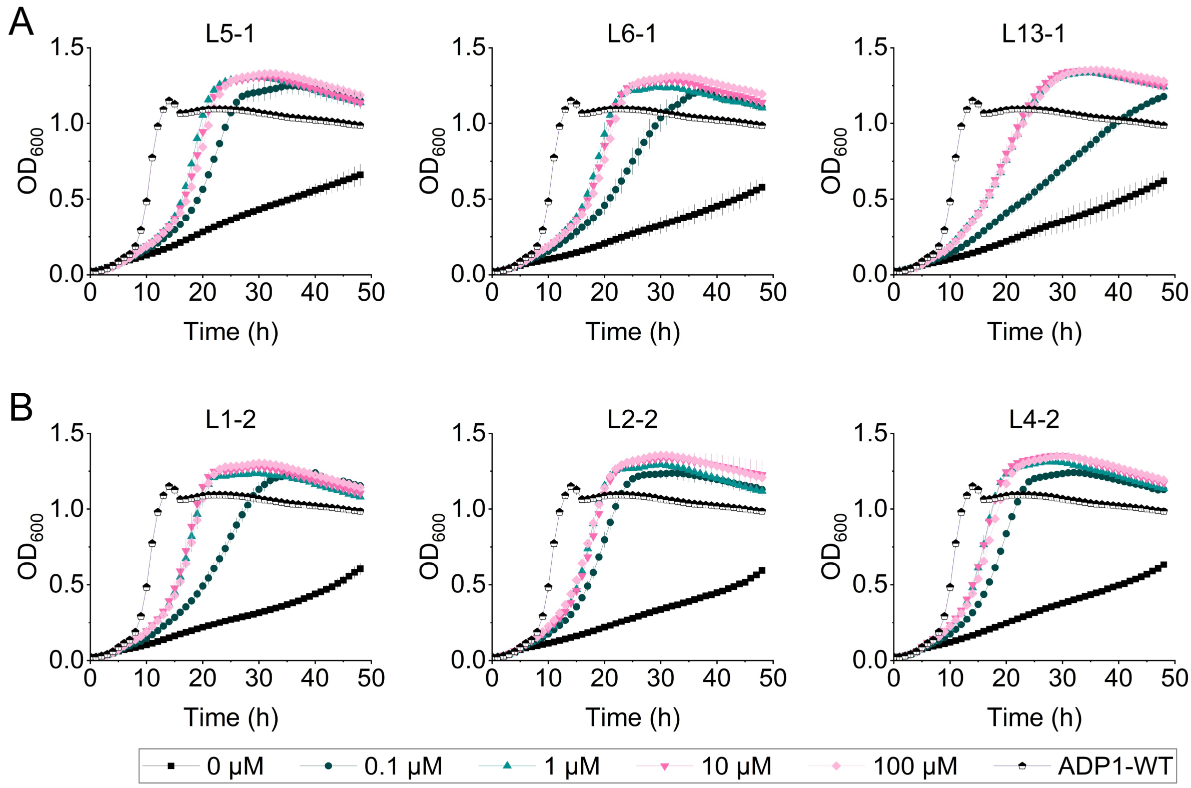


Supplementary Figure S13. Growth profile of L strains exposed to varying AHL concentrations (0–100 µM) over 48 h. Growth of (A) $ADP1_{SYNPCA}$-1 and (B) $ADP1_{SYNPCA}$-2 was monitored in MSM containing 25 mM PCA. ADP1-WT strain was used as a control. Data points represent mean values ± standard deviations from three biological replicates (n = 3).

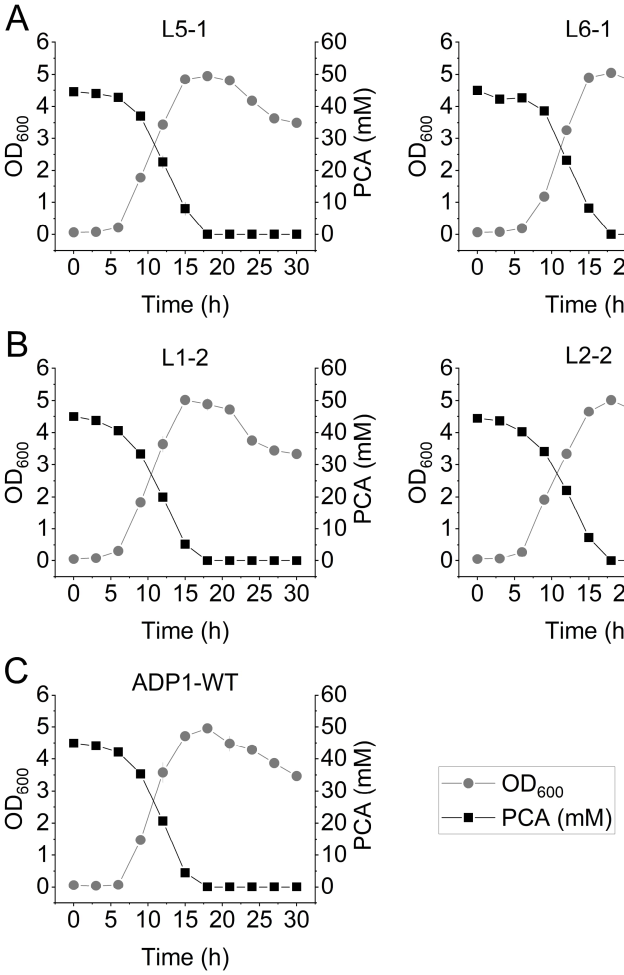


Supplementary Figure S14. Shake flask cultures of (A) ADP1$_{SYNPCA}$-1 L5, L6, L13, (B) ADP1$_{SYNPCA}$-2 L1, L2, L4, and (C) ADP1-WT in MSM with 50 mM PCA. Cell growth, dark grey circles; PCA, black squares. Data points represent the means ± standard deviations calculated from three biological replicates (n=3).

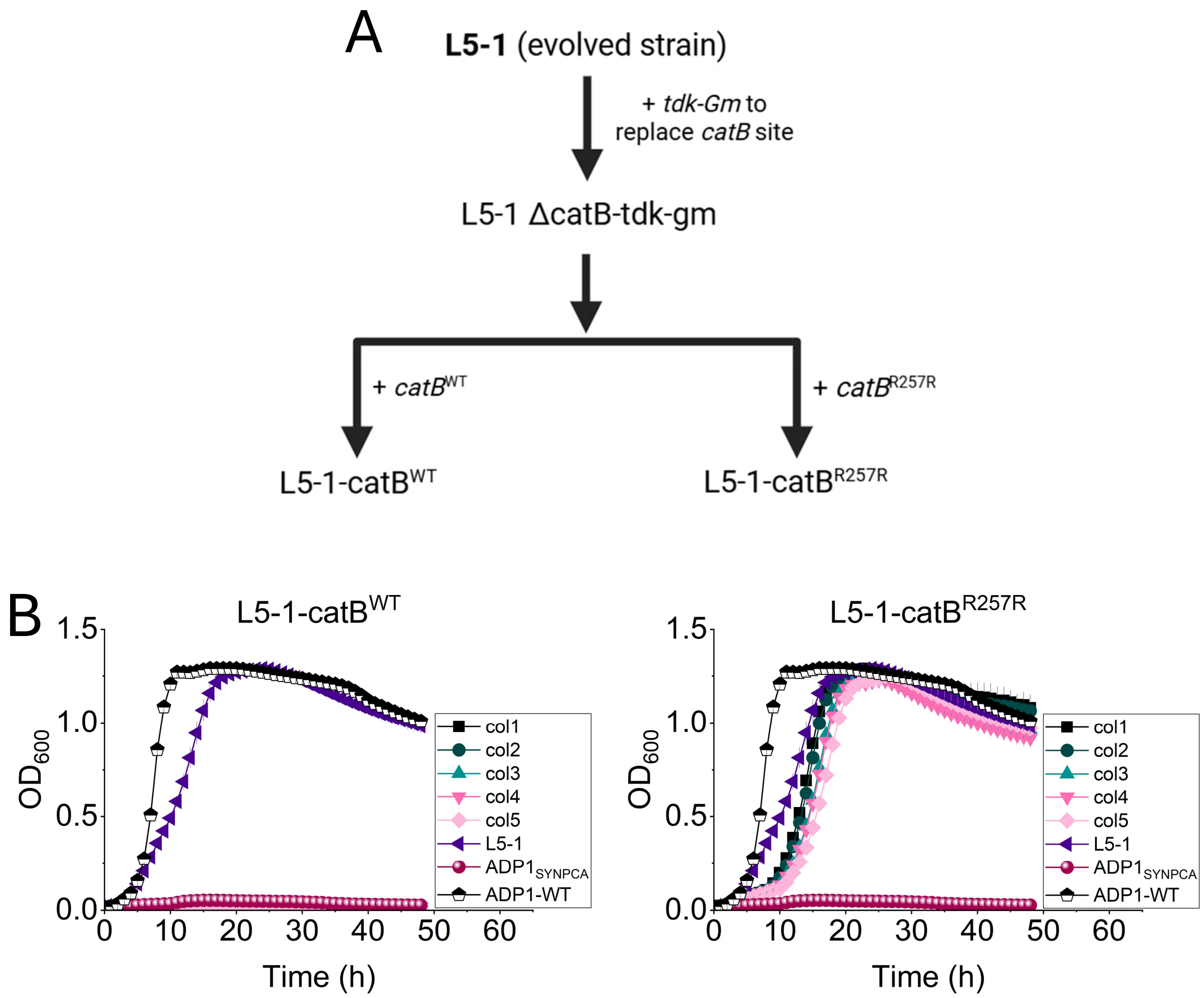


Supplementary Figure S15. Evaluation of catB$^{WT}$ and catB$^{R257R}$ in L5-1 strain. A) Construction step of L5-1 containing catB$^{WT}$ or catB$^{R257R}$, B) Growth curves of L5-1 carrying catB$^{WT}$ and catB$^{R257R}$ from five independent colonies per strain (col1 - col5). Parental strain (ADP1$_{SYNPCA}$), L5-1, L5-1-catB$^{WT}$, L5-1-catB$^{R257R}$, and ADP1-WT were cultivated in MSM containing 25 mM PCA. The synthetic pathway expression was induced with 10 µM AHL. Optical density at 600 nm ($OD_{600}$) was measured every 1 h in Tecan Spark (Switzerland) for 48h. Data points represent the mean ± standard deviation from three biological replicates (*n* = 3).

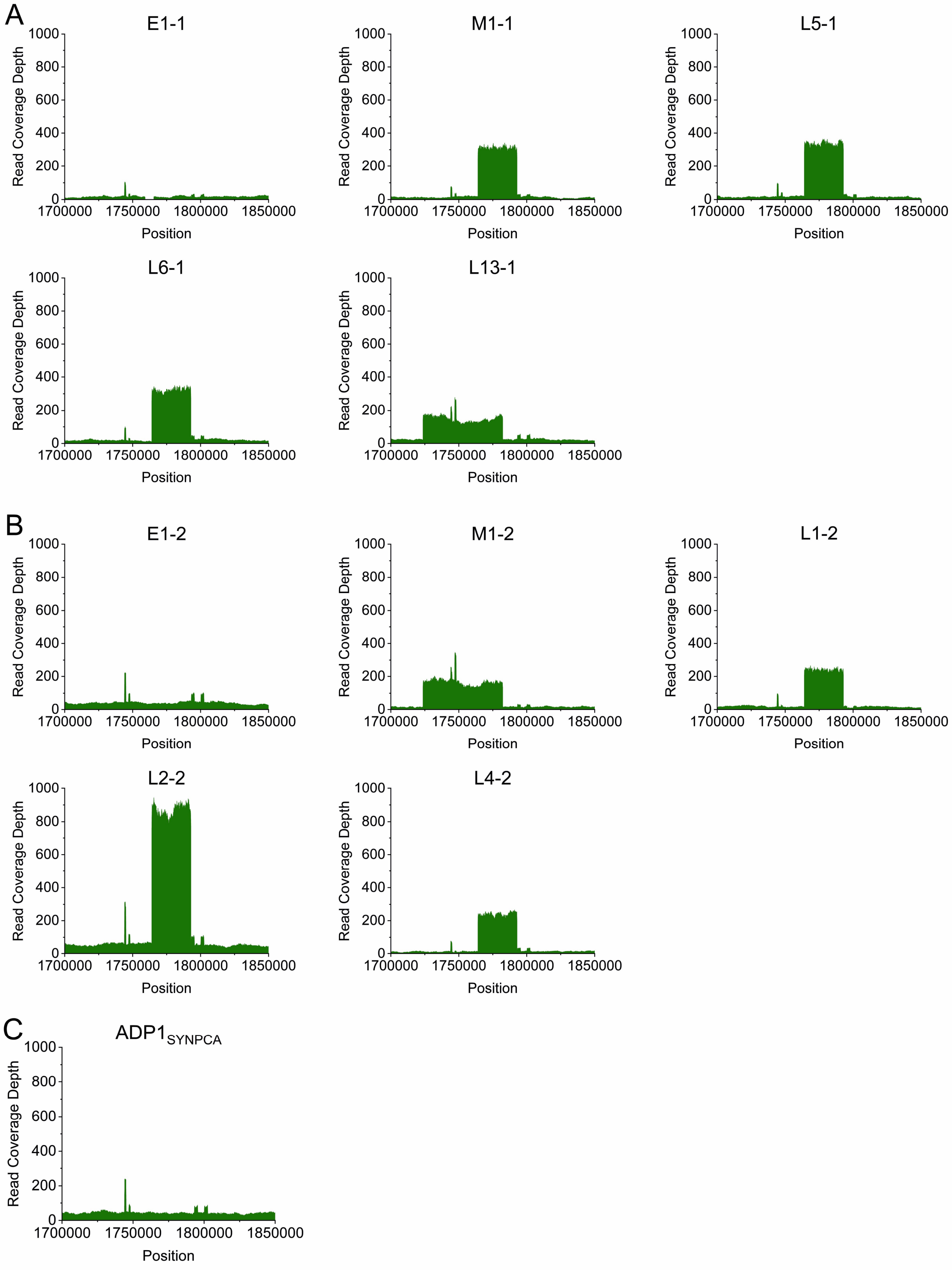


Supplementary Figure S16. Read depth coverage region of ACIAD1814-ACIAD1845 (~29 kb) (M1-1, L5-1, L6-1, L1-2, and L2-2) and ACIAD1774-ACIA1829 (~58 kb) (L13-1 and M1-2) of (A, B) evolved strains compared to (C) parental strain ($ADP1_{SYNPCA}$). Details of multiplied regions are available in Supplementary Table S9. Multiplied regions were detected only in mid and late isolates, with region ACIAD1814-ACIAD1845 more frequently observed in evolved strains.

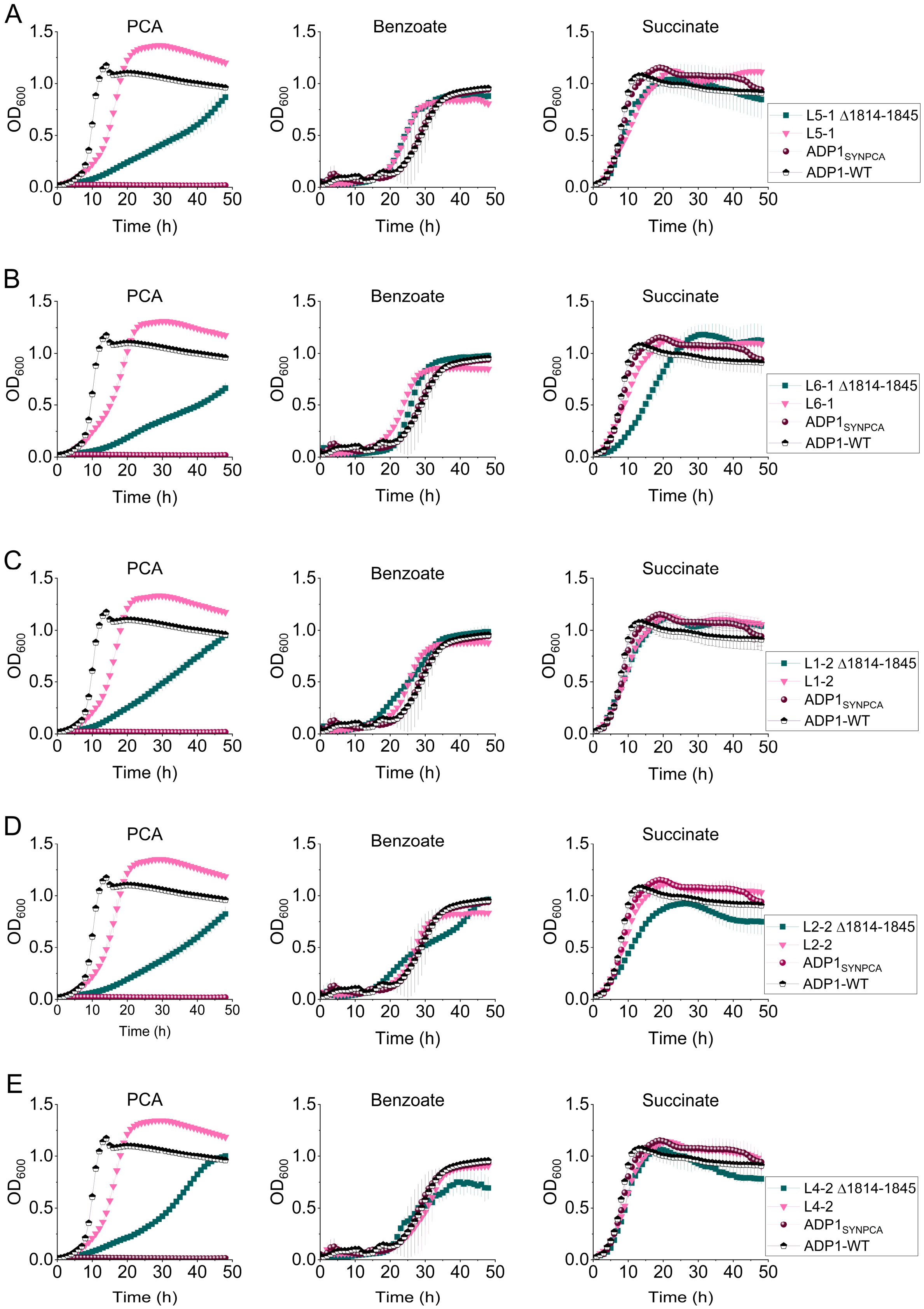


Supplementary Figure S17. Growth curves of deletion strains lacking multiplied regions (ACIAD1814-1845). ADP1, parental, evolved (L5-1, L6-1, L1-2, L2-2, and L4-2), and deletion strains (Δ1814-1845) were cultivated in minimal medium containing different carbon sources:

25 mM PCA, 10 mM benzoate, or 50 mM succinate. Optical density at 600 nm ($OD_{600}$) was measured every 1 h in Tecan Spark (Switzerland) for 48h. Data points represent the mean ± standard deviation from three biological replicates (*n* = 3).

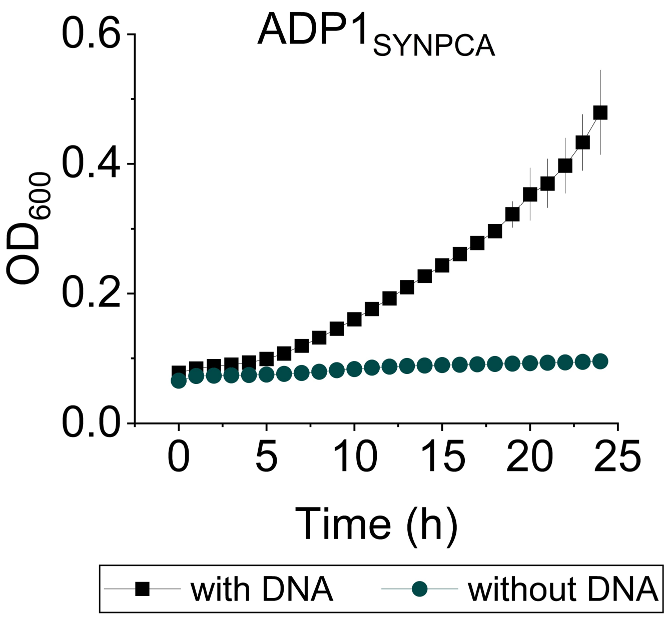


Supplementary Figure S18. Reverse engineering of $ADP1_{SYNPCA}$ in MSM with 25 mM PCA. Cells were pre-incubated in ABMS for 6 h with or without DNA, centrifuged, and transferred to fresh MSM containing 25 mM PCA for overnight cultivation. Cultures were then diluted (10 µL into 190 µL) into 96-well plates and monitored for 24 h. Wells showing growth were streaked on LB agar supplemented with Km to obtain single colonies. Data points represent the mean ± standard deviation from six technical replicates (*n* = 6).

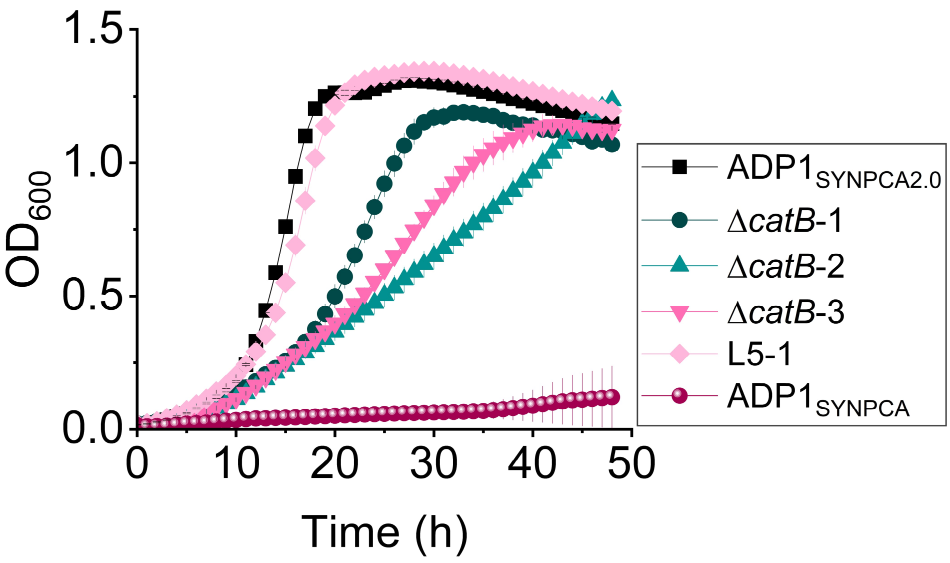


Supplementary Figure S19. Growth profile of $ADP1_{SYNPCA}$ Δ*catB* strains. Parental ($ADP1_{SYNPCA}$), evolved (L5-1), $ADP1_{SYNPCA2.0}$, and three colonies of $ADP1_{SYNPCA}$ *catB* deletion strains (Δ*catB*-1 – Δ*catB*-3) were cultivated in minimal medium containing 25 mM PCA as the sole carbon source. Optical density at 600 nm ($OD_{600}$) was measured every 1 h in Tecan Spark (Switzerland). Data points represent the mean ± standard deviation from three biological replicates (*n* = 3).

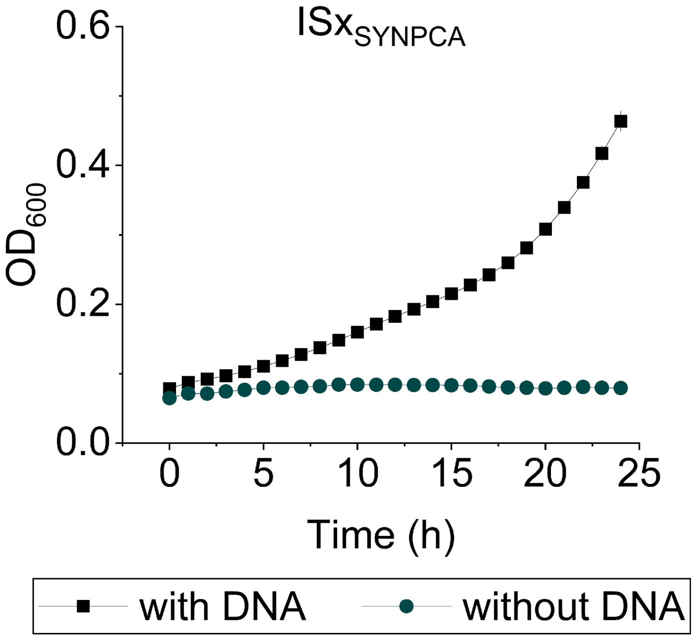


Supplementary Figure S20. Reverse engineering of $ISx_{SYNPCA}$ in MSM with 25 mM PCA. Cells were pre-incubated in ABMS for 6 h with or without DNA, centrifuged, and transferred to fresh MSM containing 25 mM PCA for overnight cultivation. Cultures were then diluted (10 µL into 190 µL) into 96-well plates and monitored for 24 h. Wells showing growth were streaked on LB agar supplemented with Km to obtain single colonies. Data points represent the mean ± standard deviation from six technical replicates (*n* = 6).

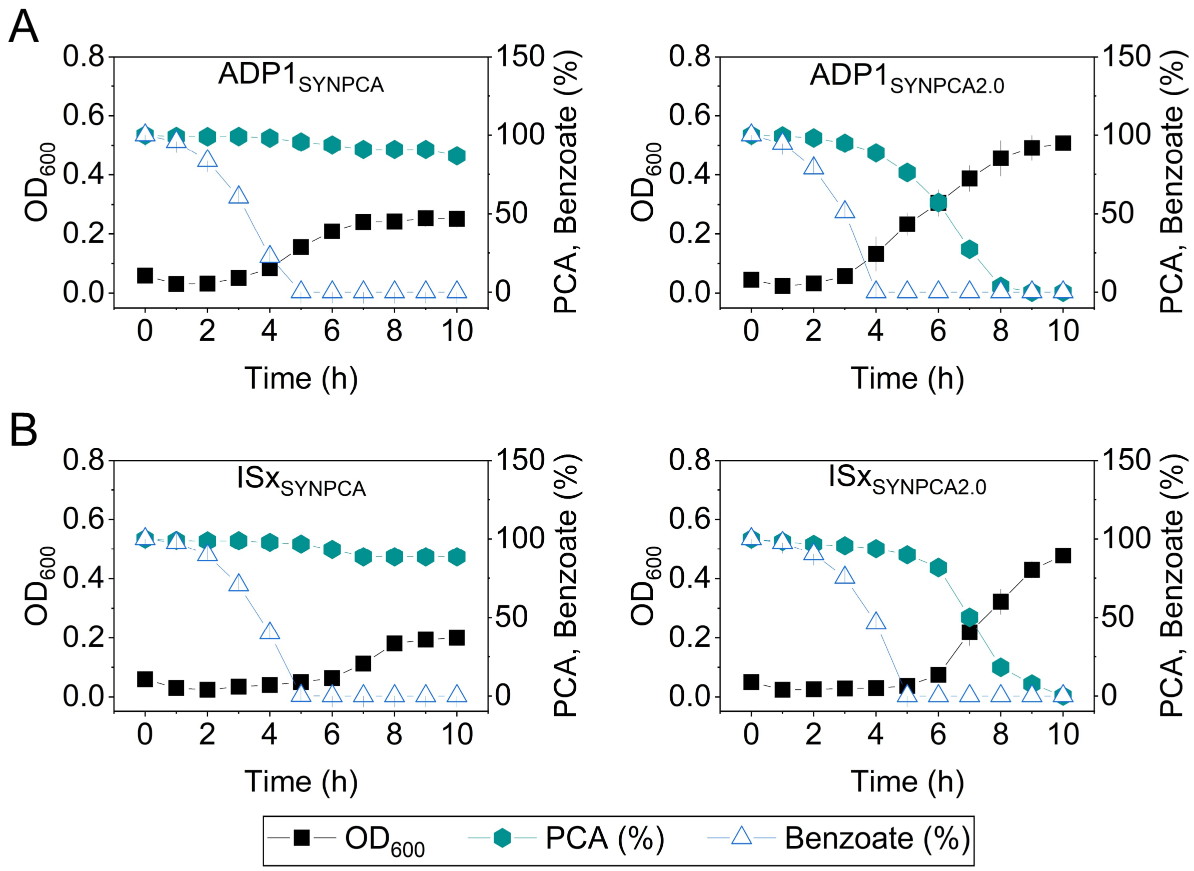


Supplementary Figure S21. Mixed-substrates consumption in ADP1 engineered strain. Batch cultures of A) $ADP1_{SYNPCA}$ and $ADP1_{SYNPCA2.0}$, B) $ISx_{SYNPCA}$ and $ISx_{SYNPCA2.0}$ strain were grown in MSM containing 2 mM benzoate and 2 mM PCA as carbon sources. Substrate concentrations were measured by HPLC at indicated time points. While $ADP1_{SYNPCA}$ and $ISx_{SYNPCA}$ exhibited growth supported solely by benzoate consumption, the strain containing *catB* mutation ($ADP1_{SYNPCA2.0}$ and $ISx_{SYNPCA2.0}$) displayed co-utilization of both benzoate and PCA. Data represents the mean ± SD of three biological replicates (n=3).

### References


1. Suárez,G.A., Dugan,K.R., Renda,B.A., Leonard,S.P., Gangavarapu,L.S. and Barrick,J.E. (2020) Rapid and assured genetic engineering methods applied to Acinetobacter baylyi ADP1 genome streamlining. *Nucleic Acids Research*, 48, 4585–4600.
